# On-target toxicity limits the efficacy of CDK11 inhibition against cancers with 1p36 deletions

**DOI:** 10.1101/2025.08.03.668359

**Authors:** Linda Julian, Lisa Crozier, Devon Lukow, Sanat Mishra, Aditi Swamy, Ryan A. Hagenson, Peter Sennhenn, Erin L. Sausville, Brianna Mendelson, Claudio Chuaqui, Lu Qiao, Anand Vasudevan, Kuan-Ting Lin, Sonam Bhatia, Thierry Bertomeu, Andrew Chatr-aryamontri, Li Zhang, Matthew G. Rees, Melissa M. Ronan, Jennifer A. Roth, Timothy Nottoli, Suxia Bai, Jayalakshmi Lakshmipathi, Viswanathan Muthusamy, Jonathan G. Van Vranken, Steven P. Gygi, Sarah L. Thompson, Joan C. Smith, Kendall Anderson, Sanjana Shah, Ranjit S. Bindra, Martin Akerman, David L. Spector, Adrian R. Krainer, Jason M. Sheltzer

**Affiliations:** Yale University, 300 George Street, New Haven, CT 06511; Cold Spring Harbor Laboratory, 1 Bungtown Road, Cold Spring Harbor, NY 11724; Trans MedChem Consulting, Franz-Joseph-Str. 7 D-80801, München, Germany; Meliora Therapeutics, 238 Main Street, Cambridge, MA 02142; Institute for Research in Immunology and Cancer, ChemoGenix platform, Université de Montréal, 2950 Chemin de Polytechnique, Montréal, QC H3T-1J4, Canada; Broad Institute of MIT and Harvard, 415 Main Street, Cambridge, MA 02142; Department of Comparative Medicine, Yale University, 310 Cedar Street, New Haven, CT 06520; Harvard Medical School, 240 Longwood Avenue, Boston, MA 02115; Envisagenics, 30-02 48th Ave, Long Island City, NY 11101

## Abstract

The cyclin-dependent kinase CDK11 is an understudied kinase that has been the subject of conflicting reports regarding its function in cancer. Here, we combine genetic and pharmacological approaches to demonstrate that CDK11 is a critical regulator of cancer cell survival that is required for RNA splicing and the expression of homologous recombination genes. Inhibition of CDK11 disrupts genome stability, promotes the retention of intronic sequences in mature mRNAs, and induces synthetic lethality with PARP inhibitors. Through integrative analysis of functional genomics datasets, we identify heterozygous deletions of chromosome 1p36 - which encompasses CDK11 and its activating cyclin CCNL2 - as a recurrent and predictive biomarker of sensitivity to CDK11 inhibition. To assess the therapeutic potential of CDK11, we develop MEL-495R, a selective and orally bioavailable CDK11 inhibitor. Additionally, we establish a genetically-engineered mouse model that allows us to differentiate between the on-target and off-target effects of CDK11 inhibitors *in vivo.* Using this platform, we demonstrate that MEL-495R induces widespread on-target toxicity, revealing a narrow therapeutic index. Together, these findings define CDK11 as a core cancer dependency, uncover a chromosomal deletion that sensitizes tumors to CDK11 inhibition, and provide a generalizable strategy for deconvolving drug efficacy and toxicity *in vivo* for novel oncology targets.

## INTRODUCTION

Cyclin-dependent kinases (CDKs) are the central drivers of the mitotic cell cycle and serve as signaling hubs that regulate multiple aspects of eukaryotic biology^1,2^. Due to their key roles in cell division, CDKs are also promising targets for cancer therapy^3,4^. The CDK4/CDK6 inhibitors abemaciclib, palbociclib, and ribociclib have received FDA approval for use in hormone receptor-positive breast cancer, and drugs targeting several other CDKs are currently undergoing clinical trials^5^. Thus, there is substantial clinical precedent and potential for targeting the CDK family. However, the human genome encodes 20 different CDKs, and the high degree of homology between them makes it challenging to develop inhibitors that are specific for individual members of this protein family. Additionally, the cellular consequences of individually targeting most CDKs remain unknown.

CDK11 is a poorly characterized member of the CDK family (see Supplemental Text 1 for a history of the controversy and confusion regarding the naming of this kinase). As with other CDKs, CDK11 function is controlled by an activating cyclin, called Cyclin L^6^. Early experiments using RNAi identified various roles for CDK11 in mitosis and splicing, though its specific functions in these processes remain elusive^7–9^. Prior research into CDK11’s role in malignant growth has come to differing conclusions, with some publications implicating CDK11 as important for cancer cell proliferation while other reports have identified CDK11 as a tumor suppressor^10–23^. Certain promiscuous pan-CDK inhibitors show some activity against CDK11, but the lack of potent and selective inhibitors of CDK11 has slowed research in this area^24^. Currently, CDK11 is listed as an “understudied target” by the NIH’s Illuminating the Druggable Genome Consortium, underscoring the significant unmet interest in characterizing this kinase^25^.

We recently reported the discovery of the first-ever selective inhibitor of CDK11^26^. This compound, called OTS964, was initially developed as an inhibitor of the PBK kinase^27^. However, we found that CRISPR-mediated deletion of PBK did not alter cellular sensitivity to OTS964, indicating that this compound’s cytotoxic effects are mediated by some other protein or proteins. We derived cancer cell lines that were resistant to OTS964, and we determined that every drug-resistant clone harbored a mutation in the kinase domain of CDK11. Introducing these mutations into drug-naïve cells was sufficient to confer resistance to OTS964, implicating this drug as a CDK11 inhibitor. Subsequently, the co-crystal structure of OTS964 bound to CDK11 was solved, verifying that OTS964 is a Type I ATP-competitive inhibitor and shedding light on the structural determinants of its specificity^28^.

The development of therapeutic interventions against novel oncology targets like CDK11 is often stymied by pervasive challenges evaluating drug toxicity. As cancer drugs frequently impact fundamental cellular processes, the identification of a therapeutic window allowing safe and effective dosing is crucial. Moreover, when a drug designed against a novel target does exhibit toxicity, it is difficult to determine whether the observed side effects result from the inhibition of its intended target in normal tissue (on-target toxicity) or from unintended interactions with other proteins (off-target toxicity). This distinction is crucial for assessing if a drug’s adverse effects can be mitigated through structural optimization or whether target suppression is inherently harmful.

One approach to evaluate the consequences of inhibiting a novel cancer target is to use inducible RNAi or another genetic intervention to ablate a gene-of-interest in adult mice^29,30^. This allows a careful determination of the effects of target suppression on mammalian physiology. However, as this approach uses a genetic rather than pharmacological intervention, it may fail to accurately recapitulate drug-based inhibition in terms of kinetics, tissue distribution, and completeness of target suppression. An ideal system for toxicity evaluation would allow the actual drug candidate itself to be tested directly, as this would provide a more clinically-relevant evaluation of the consequences of target suppression and of the drug’s safety profile. An experimental approach to rigorously differentiate between on-target and off-target toxicity *in vivo* for inhibitors of novel cancer targets has not been described.

In this manuscript, we apply complementary genetic and pharmacological approaches to characterize the consequences of CDK11 inhibition. We report the discovery of a common genomic alteration in tumors that confers enhanced sensitivity to CDK11 inhibition across diverse cancer types. Additionally, we develop a genetically-engineered mouse model that allows us to definitively differentiate between the on-target and off-target effects of CDK11 inhibitors *in vivo*. This approach provides a broadly-applicable strategy for assessing compound selectivity *in vivo* and offers critical insight into the therapeutic potential - and limitations - of targeting CDK11 for cancer therapy.

## RESULTS

### OTS964 is a selective inhibitor of CDK11

In humans, the CDK11 protein is encoded by two closely-related paralogs, *CDK11A* and *CDK11B*^31,32^. Additionally, multiple isoforms of CDK11 have been described, including the p58 isoform (which encodes only the C-terminal kinase domain) and the p110 isoform (which encodes the full-length protein)(Fig. 1A). We previously identified a point mutation in the kinase domain of CDK11, CDK11B^G579S^, that confers resistance to OTS964^26^. To further characterize the relationship between OTS964 and CDK11, we assessed the ability of OTS964 to bind to CDK11 *in cellulo*. We performed NanoBRET target engagement assays, which quantify the ability of a drug of interest to prevent the binding between a fluorescent tracer and a luciferase-tagged protein^24^. We found that OTS964 exhibits potent binding to CDK11A and CDK11B at concentrations below 100 nM (Fig. S1A). In contrast, the CDK4/6 inhibitor palbociclib exhibits minimal binding to either protein. Next, we assessed OTS964 binding across the kinome by performing a KiNativ profiling assay, which measures the ability of a compound to block the accessibility of kinase domains in native cellular lysate^33^. We found that treatment with 100 nM OTS964 resulted in 72% inhibition of CDK11A and CDK11B (which cannot be distinguished in this assay) and did not significantly inhibit any other cellular kinases (Fig. S1B and Table S1).

**Figure 1.**
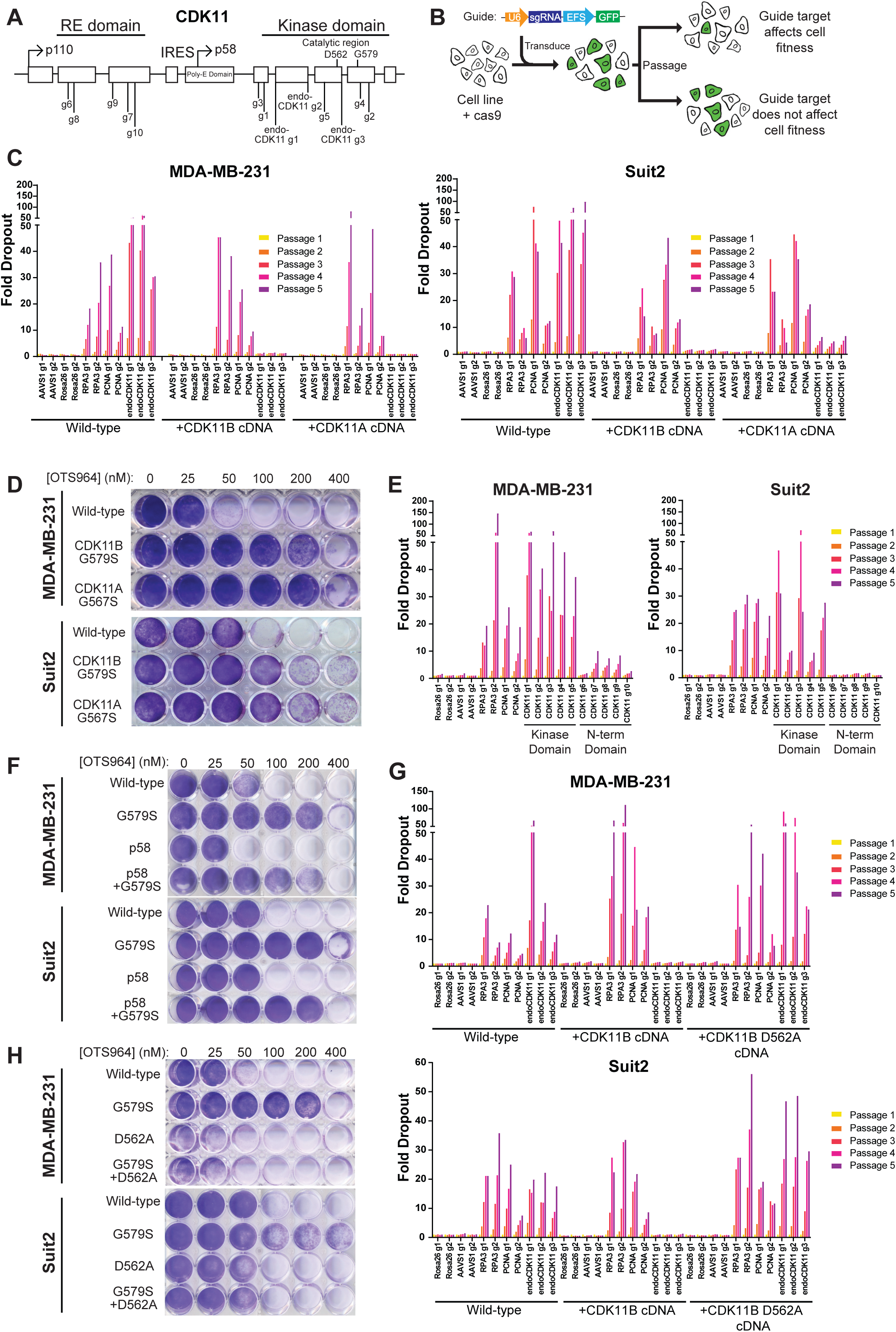
Genetic and pharmacological inhibition reveal that CDK11 kinase activity is required for cancer cell viability. A) Schematic of the CDK11 locus. The relative locations of the sequences targeted by gRNAs used in the dropout experiments are shown. B) Schematic of the CRISPR-based dropout assay for assessing cancer cell viability^35^. C) Dropout assays in MDA-MB-231 (left) and Suit2 (right) cell lines. Plots display the fold dropout of cells transduced with guides targeting negative control genes (AAVS1, Rosa26), positive control genes (RPA3, PCNA), or endogenous CDK11A and CDK11B (endoCDK11). Dropouts were performed in wild-type cells and cells transduced with vectors expressing gRNA-resistant CDK11A or CDK11B cDNA. Complete dropout results and statistical testing are included in Table S2A. D) Crystal violet staining of MDA-MB-231 (top) and Suit2 (bottom) cells treated with OTS964 in the presence of cDNAs expressing either CDK11B^G579S^ or CDK11A^G567S^. E) Dropout assays in MDA-MB-231 (left) and Suit2 (right) cell lines. Plots display the fold dropout of cells transduced with guides targeting negative control genes (Rosa26, AAVS1), positive control genes (RPA3, PCNA), the CDK11 kinase domain, or the CDK11 N-terminal domain. Complete dropout results and statistical testing are included in Table S2B. F) Crystal violet staining of MDA-MB-231 (top) and Suit2 (bottom) cells treated with OTS964 in the presence of cDNAs expressing either full-length CDK11B^G579S^, the CDK11B-p58 isoform, or the CDK11B-p58 isoform with the G579S mutation. G) Dropout assays in MDA-MB-231 (top) and Suit2 (bottom) cell lines. Plots display the fold dropout of cells transduced with guides targeting negative control genes (Rosa26, AAVS1), positive control genes (RPA3, PCNA), or endogenous CDK11. Dropouts were performed in wild-type cells and in cells expressing gRNA-resistant CDK11B or gRNA-resistant, kinase dead CDK11B cDNA (D562A). Complete dropout results and statistical testing are included in Table S2C. H) Crystal violet staining of MDA-MB-231 (top) and Suit2 (bottom) cells treated with OTS964 in the presence of cDNAs expressing either CDK11B^G579S^, kinase-dead CDK11B (D562A), or the kinase-dead G579S double mutant (D562A/G579S).

We subsequently tested the ability of OTS964 to inhibit various members of the CDK family in *in vitro* kinase assays. We found that OTS964 inhibits CDK11B function with an IC50 of 49 nM, which was consistent with the potencies that we observed *in cellulo* and in cell lysate binding assays (Fig. S1C). In contrast, palbociclib did not inhibit CDK11B activity at concentrations up to 10 µM. Additionally, OTS964 exhibited reduced or no activity against 14 other CDKs (Fig. S1D). After CDK11, the next most potent target of OTS964 was CDK9, which exhibited a 13-fold higher IC50 value (650 nM) compared to CDK11B. Finally, we assessed the ability of OTS964 to inhibit CDK11 function *in cellulo*. We found that 100 nM OTS964 decreased the phosphorylation of Ser2 on RNAPII, which is a previously-described target of CDK11 (Fig. S1E)^34^. Expression of the CDK11B^G579S^ resistance mutation restored Ser2 phosphorylation in the presence of OTS964, demonstrating that the decrease in phosphorylation is an on-target effect of CDK11 inhibition. In total, these results indicate that OTS964 is a potent and selective inhibitor of CDK11 function *in vitro* and *in cellulo*.

### CDK11 kinase activity is required for cancer cell viability

Prior studies investigating CDK11’s role in cancer have yielded conflicting conclusions. While some research has suggested that CDK11 is essential for cancer cell growth, others have reported that it acts as a tumor suppressor, potentially inhibiting rather than promoting cellular proliferation^10–23^. Moreover, the functional importance of CDK11’s multiple paralogs and isoforms remains largely unexplored. To clarify the role of CDK11 in cancer cell fitness, we performed CRISPR competition assays in a breast cancer cell line (MDA-MB-231) and a pancreas cancer cell line (Suit2). In these assays, cells were transduced at a low multiplicity-of-infection with gRNAs targeting a gene of interest in a vector that co-expresses GFP. The loss of GFP-expressing cells over time indicates that the target of the gRNA is essential for cellular fitness (Fig. 1B). As negative controls for this assay, we used gRNAs that target the non-essential loci *Rosa26* and *AAVS1*, and as positive controls, we used gRNAs targeting the genes that encode the pan-essential replication factors *RPA3* and *PCNA*^26,35^.

We first designed gRNAs that recognize sequences in the kinase domain that are shared by both *CDK11A* and *CDK11B*. We found that gRNAs targeting these loci exhibited significant dropout in both MDA-MB-231 and Suit2 cells, comparable to the degree of dropout exhibited by gRNAs targeting *RPA3* and *PCNA* (Fig. 1C and Table S2). Next, we repeated these competition experiments in cells transduced with gRNA-resistant *CDK11A* or *CDK11B* cDNA. We found that the expression of either cDNA was sufficient to prevent the dropout of CDK11-targeting gRNAs. These results demonstrate that the lethality of the CDK11-targeting gRNAs is due to the on-target ablation of CDK11, and the expression of either *CDK11A* or *CDK11B* is generally sufficient to support cellular viability. To further explore the redundancy between *CDK11A* and *CDK11B*, we established MDA-MB-231 and Suit2 cells that express either CDK11B^G579S^ or the homologous mutation in CDK11A, CDK11A^G567S^. We found that the expression of either protein was sufficient to restore the viability of cancer cells treated with a lethal concentration of OTS964, verifying the results of our CRISPR experiment and indicating that the expression of either CDK11 paralog is generally sufficient for cellular viability (Fig. 1D).

Next, we explored whether the expression of the full-length CDK11^p110^ isoform is required for viability. As above, we performed complementary experiments using both CRISPR and OTS964 to perturb CDK11 function. First, we conducted CRISPR competition assays using a panel of ten gRNAs that targeted either the CDK11 kinase domain or the N-terminal domain, the latter of which is included in CDK11^p110^ but not CDK11^p58^. We found that gRNAs targeting the kinase domain exhibited significantly greater dropout than gRNAs targeting the N-terminal domain, indicating that the kinase domain likely encodes the key function(s) of the protein that are required for viability (Fig. 1E). Next, we generated MDA-MB-231 and Suit2 cells that express the G579S resistance mutation in the truncated CDK11^p58^ protein. We found that the expression of this isoform was sufficient to restore the viability of cells treated with a lethal concentration of OTS964 (Fig. 1F). In total, these results demonstrate that the expression of the CDK11 kinase domain is sufficient for viability in these cancer cell lines.

Finally, we sought to determine whether CDK11’s kinase activity itself is required for cellular viability. We performed CRISPR competition assays in cells expressing either gRNA-resistant *CDK11B* cDNA or *CDK11B* cDNA encoding a kinase-inactivating mutation, D562A. We found that the expression of kinase-dead *CDK11B* cDNA was incapable of rescuing the lethality caused by CDK11-targeting gRNAs (Fig. 1G). Similarly, while the expression of CDK11B^G579S^ was sufficient to block the effects of OTS964, the expression of the CDK11B^D562A,G579S^ double-mutant had no effect on OTS964 sensitivity (Fig. 1H). These experiments provide genetic and pharmacological evidence that CDK11’s kinase activity is required for cancer cell viability.

### CDK11 is required for accurate splicing

To explore CDK11’s role in cancer, we treated four cancer cell lines expressing either wild-type CDK11B or CDK11B^G579S^ with OTS964 and then analyzed changes in gene expression (Table S3). Consistent with CDK11’s role in splicing^7,8,36^, we found that CDK11 inhibition results in pervasive alterations in splicing patterns throughout the transcriptome, including a notable increase in intron retention events (Fig. 2A and Table S4). While read-depth analysis of untreated cells revealed clear patterns indicative of intron excision, we found that introns were retained in mature mRNA transcripts across hundreds of genes in OTS964-treated cells (Fig. 2A-2B). Using PCR primers designed to amplify across either exon-exon or intron-exon junctions, we confirmed that OTS964 treatment results in a ∼30-fold increase in intron retention in *CCNA2* transcripts and a ∼15-fold increase in intron retention in *CDK9* transcripts (Fig. S2A-B). The expression of CDK11B^G579S^ rescued wild-type splicing patterns in the presence of OTS964, demonstrating that these alterations are an on-target consequence of CDK11 inhibition (Fig. 2A-B and S2B). Gene Set Enrichment Analysis (GSEA) revealed that transcripts with retained introns were primarily associated with gene expression, splicing, and cell cycle progression (Fig. 2C and Table S5).

**Figure 2.**
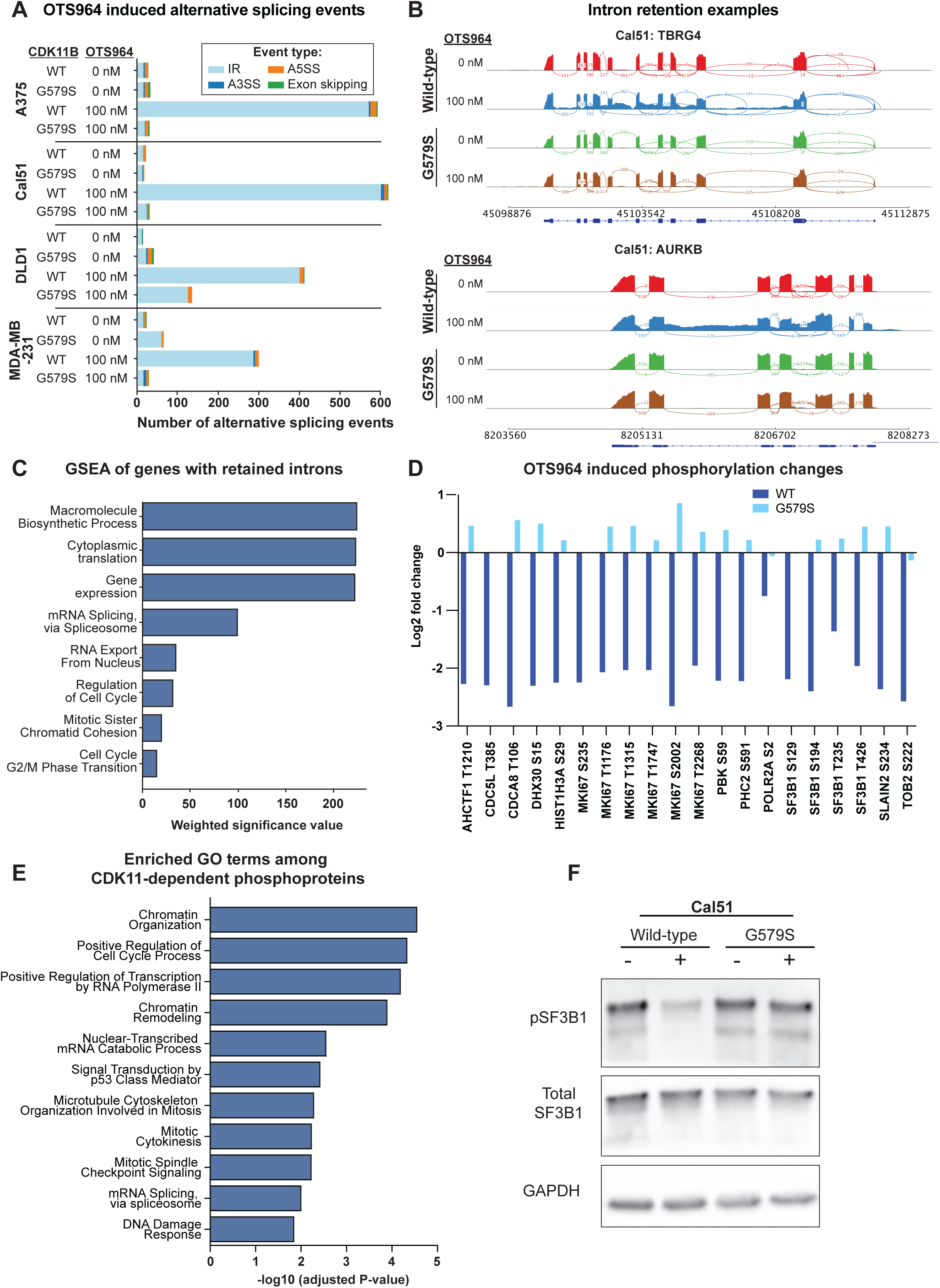
CDK11 inhibition interferes with splicing and promotes the retention of intronic sequences. A) Bar graph showing alternate splicing events in A375, Cal51, DLD1, and MDA-MB-231 cell lines in cells expressing wild-type CDK11B or CDK11B^G579S^ treated with 100 nM OTS964 for 6 hours. Alternative splicing events were detected using PSI-Sigma, with a delta-psi cutoff of 20 and p-value cutoff of 0.01^74^. IR: intron retention; A3SS: Alternative 3’ splice site; A5SS: Alternative 5’ splice site. Complete results are included in Table S4. B) Sashimi plots displaying the effects of OTS964 on *TBRG4* and *AURKB*. C) GSEA of transcripts with retained introns following OTS964 treatment. D) Bar graph showing phosphorylation changes in proteins associated with mitosis, gene expression, and splicing in A375 cells expressing wild-type CDK11B or CDK11B^G579S^ treated with 100 nM OTS964. Complete results are included in Table S6A. E) GO term enrichment analysis of proteins whose phosphorylation decreased after OTS964 treatment in wild-type cells but were not affected by OTS964 treatment in cells expressing CDK11B^G579S^. Complete results are included in Table S6B. F) Western blot assessing phosphorylation status of SF3B1 after treatment with DMSO or OTS964. GAPDH levels were examined as a loading control.

To uncover the cause(s) of these altered splicing patterns, we performed quantitative proteomic and phosphoproteomic analysis on cells treated with OTS964. Consistent with our results in Fig. S1E, unbiased mass spectrometry analysis confirmed a decrease in RNAPII phosphorylation in OTS964-treated cells (Fig. 2D and Table S6A). Additionally, we observed phosphorylation changes across multiple proteins involved in chromatin organization, mitosis, gene expression, and splicing (Fig. 2D-E and Table S6B). Notably, we found that OTS964 treatment decreased the phosphorylation of the key spliceosome component SF3B1 at multiple residues, and these changes were reversed by the expression of CDK11B^G579S^ (Fig. 2D and Table S6A). We further verified OTS964-induced dephosphorylation at the critical SF3B1-T313 residue via western blot (Fig. 2F).

Many transcripts that harbor incompletely-excised introns are retained in the nucleus and subsequently degraded^37–39^. We sought to determine whether transcripts that required CDK11-dependent processing were similarly confined to the nucleus. Surprisingly, RNA-seq analysis of the cytoplasmic fraction from OTS964-treated cells identified 298 intron-retention events, compared to 67 intron-retention events in the cytoplasm of untreated cells (Fig. S2C). We further verified via qRT-PCR that CDK11 inhibition results in an accumulation of mis-spliced *CCNA2* transcripts in the cytoplasm (Fig. S2D). In total, these results demonstrate that CDK11 inhibition results in pervasive splicing dysregulation, and we identified several candidate CDK11 targets that could mediate these effects.

### CDK11 is required for the expression of DNA repair genes

In addition to the splicing-related genes described above, our RNA-seq analysis revealed that OTS964 treatment results in a significant downregulation of genes associated with DNA repair (Fig. 3A and Table S7). Six hours of treatment with OTS964 resulted in the downregulation of *BRCA1*, *BRCA2*, *RAD51*, *EXO1*, and other genes involved in DNA repair and homologous recombination (Fig. 3B). These transcriptional alterations were an on-target effect of CDK11 inhibition, as the expression of CDK11B^G579S^ reversed the effects of OTS964 and restored wild-type levels of repair gene expression (Fig. 3B). Interestingly, this transcriptional repression of DNA repair genes was mechanistically distinct from the widespread splicing errors induced by OTS964. Very few DNA repair transcripts exhibited evidence of intron retention, and the repair gene transcripts that did not harbor retained introns were expressed at significantly lower levels than the few that did (Fig. S3).

**Figure 3.**
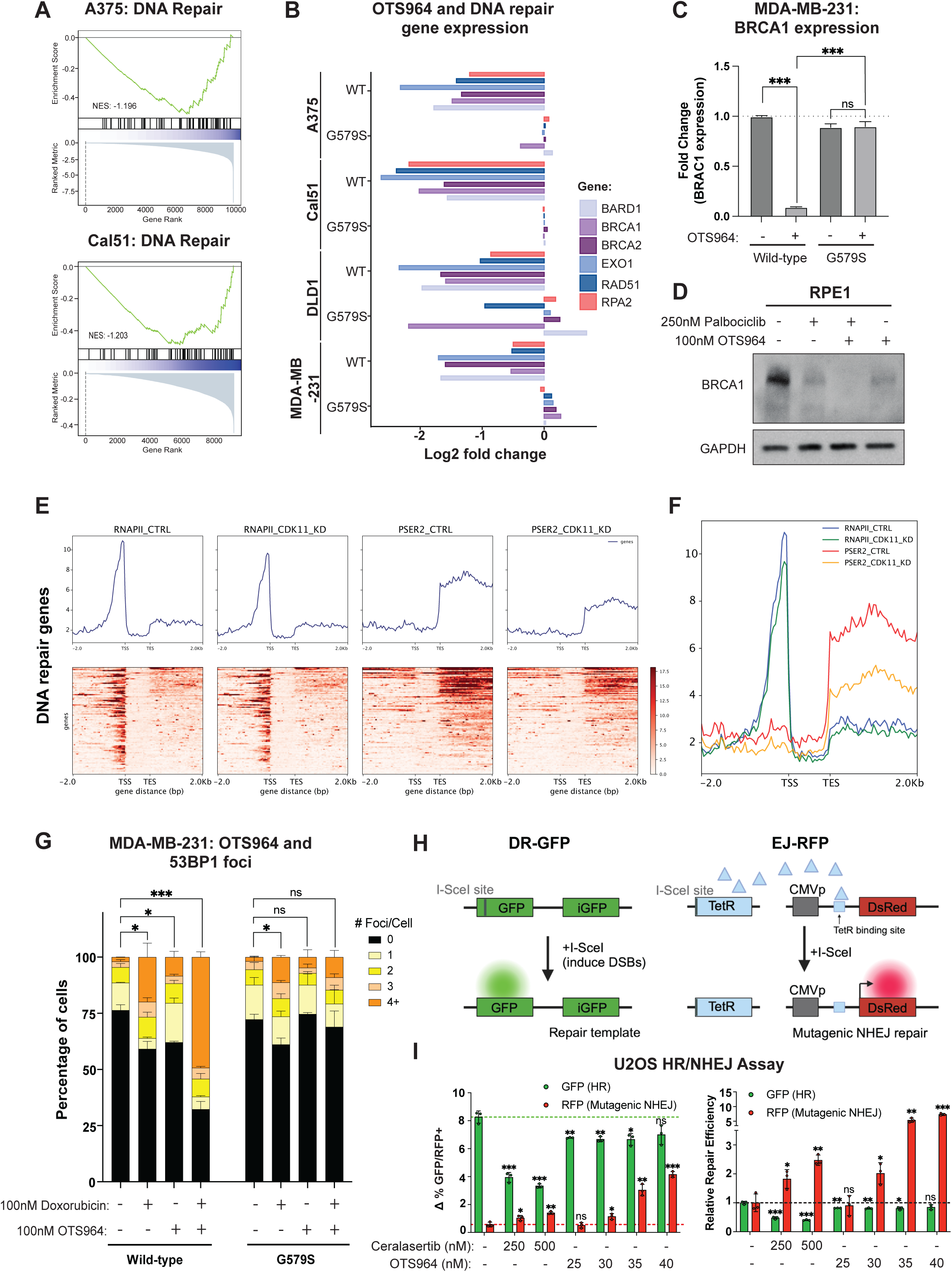
CDK11 is required for the expression of homologous recombination genes and the accurate repair of double-strand breaks. A) Gene Set Enrichment Analysis of cells treated with OTS964 reveals a downregulation of genes associated with DNA repair. Complete results are included in Table S7. B) Bar graph displaying the expression of several genes involved in homologous recombination and DNA repair in cells expressing wild-type CDK11B or CDK11B^G579S^ treated with 100 nM OTS964 for 6 hours and quantified via RNA-seq. C) Bar graph displaying the fold change in *BRCA1* expression in MDA-MB-231 cells expressing wild-type CDK11B or CDK11B^G579S^ treated with 100 nM OTS964 for 24 hours determined via qPCR. Relative gene expression levels were normalized to GAPDH. Mean ± SEM, data from three independent replicates. Statistical significance was determined using an unpaired t-test (***, P < .0005). D) Western blot assessing BRCA1 levels in RPE1 cells after treatment with DMSO, palbociclib (250nM), OTS964 (100nM), or palbociclib and OTS964 for 24 hours. GAPDH levels were examined as a control. E) ChIP–seq analyses of RNAPII and RNAPII-pSer2 occupancies on DNA repair genes, treated with control or CDK11-targeting siRNA^34^. F) Meta-plot of RNAPII and RNAPII-pSer2 occupancy across DNA repair genes. G) Bar graph showing the percentage of cells with 53BP1 foci in MDA-MB-231 cells expressing wild-type CDK11B or CDK11B^G579S^ treated with 100 nM OTS964, 100 nM doxorubicin, or OTS964 and doxorubicin. Mean ± SEM, data from three independent replicates. Statistical significance was determined using Fisher’s exact test (*, P < .05; **, P < .005; ***, P < .0005). H) Schematic of the assay used to assess the effects of OTS964 on homologous recombination and non-homologous end joining^40^. I) U2OS cells containing the HR or NHEJ plasmids described in (H) were treated with DMSO or the indicated dose of the ATR inhibitor ceralasertib or OTS964 for 48 hours. The left-hand panel shows Δ% GFP/RFP+ cells. The right-hand panel shows the relative repair efficiency of HR and NHEJ. Mean ± SEM, data from two independent replicates. Statistical significance was determined by unpaired t-tests (*, P < .05; **, P < .005; ***, P < .0005).

The homologous recombination gene *BRCA1* was one of the strongest transcripts affected by OTS964 treatment. We verified via qRT-PCR that OTS964 treatment in MDA-MB-231 cells resulted in a 10-fold downregulation of *BRCA1* transcript levels, while the expression of CDK11B^G579S^ blocked OTS964-mediated *BRCA1* repression (Fig. 3C). As *BRCA1* is a cell cycle-regulated gene, we considered the alternative possibility that the effects of OTS964 treatment on *BRCA1* expression could be an indirect consequence of CDK11’s role in cell cycle progression. To investigate this possibility, we treated cells with the CDK4/6 inhibitor palbociclib to induce a G1 arrest and then exposed the cells to OTS964. We found that co-treatment with palbociclib and OTS964 resulted in a stronger downregulation of BRCA1 expression compared to the effects of either drug alone, demonstrating that the effects of OTS964 on BRCA1 expression are not a secondary consequence of a cell-cycle arrest (Fig. 3D).

As the OTS964-driven downregulation of DNA repair genes occurs without prominent splicing alterations and independently of cell cycle dysregulation, we sought to uncover the mechanistic basis underlying the relationship between CDK11 and DNA repair gene expression. Toward that goal, we analyzed RNAPII and RNAPII-pSer2 localization by ChIP-seq in wild-type cells^34^. We found strong RNAPII-pSer2 localization over the bodies of DNA repair genes, indicative of robust transcriptional elongation (Fig. 3E). Knockdown of CDK11 decreased RNAPII-pSer2 levels at many of these genes, including *BRCA1*, *EXO1*, and *RPA2* (Fig. 3F and S4A). Genome-wide analysis of pSer2 peaks abolished by CDK11-knockdown revealed a significant enrichment of transcripts associated with DNA replication and repair (Fig. S4B and Table S8). Taken together, these data suggest that CDK11 regulates gene expression through two distinct mechanisms: ensuring accurate splicing via SF3B1, and, separately, controlling the transcription of certain genes by promoting the activation of RNAPII.

### CDK11 inhibition impairs DNA repair and results in synergistic lethality with PARP inhibitors

Our discovery that CDK11 is required for the expression of BRCA1 and other DNA repair genes raised the possibility that CDK11 inhibition could compromise homologous recombination. To investigate this hypothesis, we treated cells with OTS964, the topoisomerase poison doxorubicin, or both drugs combined, and then quantified 53BP1 foci as a marker of DNA damage. We found that OTS964 exposure increased the appearance of 53BP1 foci in both untreated and doxorubicin-treated cells (Fig. 3G). This effect was an on-target consequence of CDK11 inhibition, as the expression of the CDK11B^G579S^ mutation blocked the appearance of OTS964-induced damage foci (Fig. 3G). To directly investigate whether CDK11 is required for the repair of DNA damage, we used a DR-GFP/EJ-RFP assay, which allows the monitoring of both homologous recombination (HR)-dependent and non-homologous end-joining (NHEJ)-dependent repair of a double-strand break (Fig. 3H)^40^. As a positive control, we treated the DR-GFP/EJ-RFP cells with the ATR inhibitor ceralasertib, which resulted in a significant decrease in HR activity and a significant increase in NHEJ repair. Similarly, we found that treatment with sub-lethal doses of OTS964 caused a moderate but significant decrease in HR repair and a highly-significant increase in NHEJ (Fig. 3I).

Compromised homologous recombination causes synergistic lethality with inhibitors of PARP-dependent NHEJ^41^. We therefore sought to determine whether CDK11 inhibition exhibits a synergistic interaction when combined with PARP inhibition. We treated two triple-negative breast cancer cell lines, MDA-MB-231 and HCC1806, with OTS964 or the PARP inhibitor olaparib, either alone or in combination, and measured cell viability. In both cell lines, we found that the combination of OTS964 and olaparib resulted in greater cell killing than either drug alone (Fig. S5). Bliss analysis revealed multiple dosage combinations with synergy scores greater than 10, indicating synergistic drug interactions^42^. In total, our data demonstrate that CDK11 is required for accurate DNA repair, and inhibition of CDK11 causes synthetic lethality when combined with PARP inhibitors.

### Identification of 1p36 deletions as a biomarker for sensitivity to CDK11 inhibition

The use of a genetic biomarker to select sensitive patient populations is associated with a significant increase in clinical trial success in oncology^43^. In order to identify biomarkers capable of predicting sensitivity to CDK11 inhibition, we conducted an unbiased analysis of CRISPR screening data from the DepMap^44^. The Avana CRISPR library used to perform these screens includes three gRNAs that exhibit perfect complementarity to both *CDK11A* and *CDK11B*, and we used these gRNAs to assess each cell line’s dependence on CDK11 (Fig. S6A)^45^. We examined mutation, copy number, and transcriptional profiles of several hundred cancer cell lines and compared them to each cell line’s calculated CDK11 dependency score. We discovered that the strongest genomic biomarkers correlating with CDK11 dependency were copy number alterations affecting chromosome 1p36 (Fig. 4A and Table S9A). Cancer cell lines harboring deletions of genes encoded in the 1p36 locus were significantly more sensitive to CDK11-targeting gRNAs compared to cancer cell lines in which 1p36 was copy-neutral or amplified (Fig. 4B). Similarly, genes whose expression correlated with sensitivity to CDK11 ablation also tended to be encoded in the 1p36 locus, and low expression of these genes correlated with increasing CDK11 dependency (Fig. 4C-D and Table S9B). In contrast, we did not identify any recurrent mutations that were significantly associated with sensitivity to CDK11-targeting gRNAs (Fig. S6B and Table S9C).

**Figure 4.**
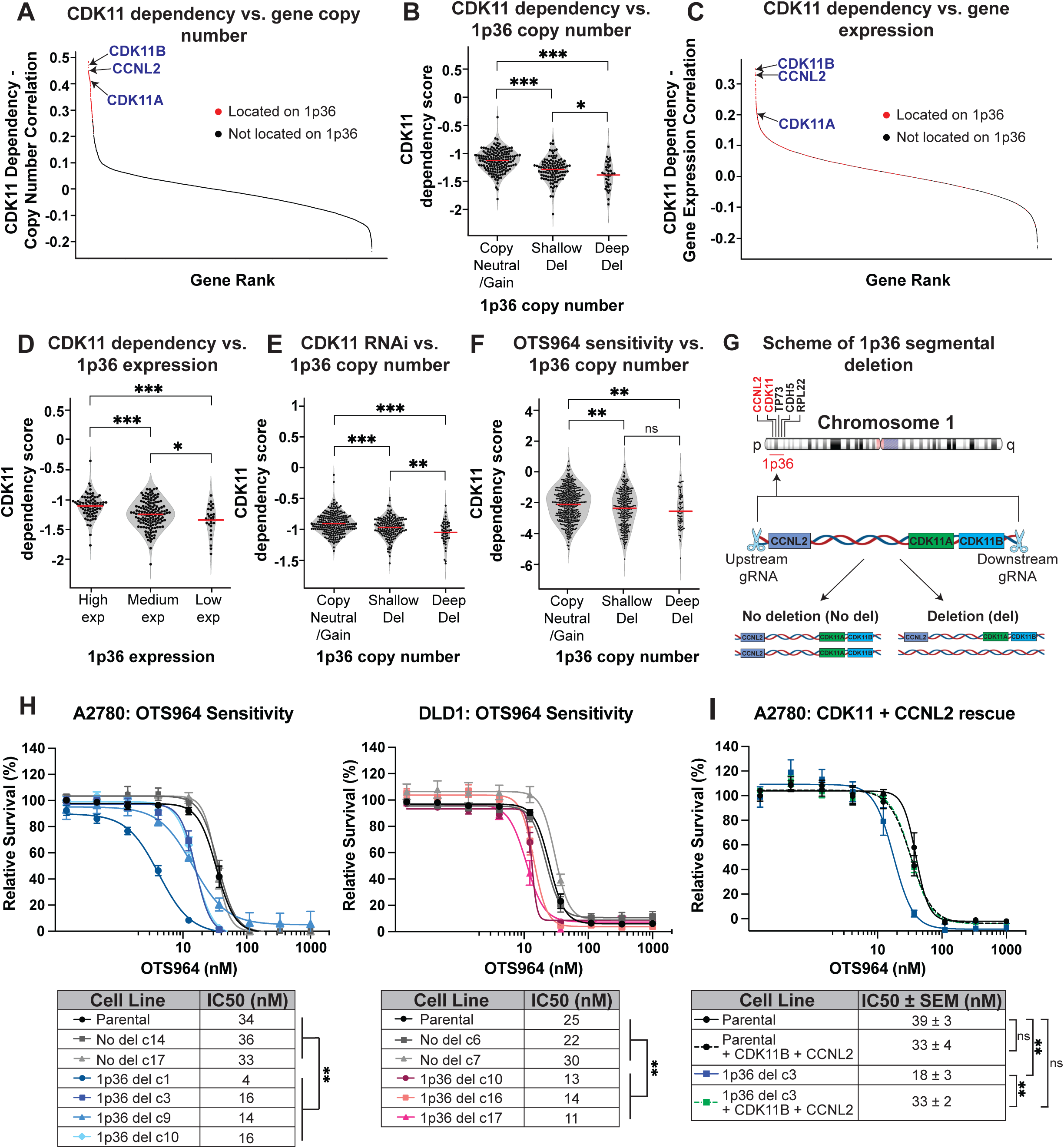
Heterozygous deletions in 1p36 confer sensitivity to CDK11 inhibition by decreasing the expression of CDK11 and CCNL2. A) Waterfall plot displaying the correlation between gene copy number and CDK11 dependency derived from CRISPR screening. Genes located on chromosome 1p36 are indicated in red. Complete results are included in Table S9A. B) Violin plots indicating the CDK11 dependency score of cancer cell lines split based on the copy number of chromosome 1p36. Every dot indicates a cancer cell line, and the red bars indicate the median of the data. C) Waterfall plot displaying the correlation between gene expression and CDK11 dependency derived from CRISPR screening. Genes located on chromosome 1p36 are indicated in red. Complete results are included in Table S9B. D) Violin plots indicating the CDK11 dependency score of cancer cell lines split based on the expression of chromosome 1p36. Every dot indicates a cancer cell line, and the red bars indicate the median of the data. E) Violin plots indicating the CDK11 dependency score of cancer cell lines from RNAi screening split based on the copy number of chromosome 1p36. Every dot indicates a cancer cell line, and the red bars indicate the median of the data. F) Violin plots indicating the sensitivity of cancer cell lines to OTS964 (123.5 nM) split based on the copy number of chromosome 1p36. Every dot indicates a cancer cell line, and the red bars indicate the median of the data. Complete results are presented in Table S10. G) A diagram of chromosome 1 indicating the position of CDK11, CCNL2, several tumor suppressors that are encoded on chromosome 1p36, and the locations targeted by gRNAs to produce 1p36 deletions using CRISPR. H) Dose-response curves showing survival of cell lines derived from (left) A2780 and (right) DLD1 treated with varying concentrations of OTS964 for 72 hours. The tested cell lines include the parental cells, cells harboring 1p36 deletions generated with CRISPR, and cells transfected with gRNAs to induce 1p36 deletions but that did not acquire a deletion of this locus. The IC50 values for each cell line are annotated in the table below. Mean ± SEM, data from three independent replicates. Statistical significance was determined by unpaired t-tests (*, P < .05; **, P < .005; ***, P < .0005). I) Dose-response curves showing survival of cell lines derived from A2780 treated with varying concentrations of OTS964 for 72 hours. The tested cell lines include the parental cells, cells harboring 1p36 deletions generated with CRISPR, and both cell types transduced with lentiviral cDNAs to overexpress both *CDK11B* and *CCNL2*. The IC50 values for each cell line are annotated in the table below. Mean ± SEM, data from three independent replicates. Statistical significance was determined by unpaired t-tests (*, P < .05; **, P < .005; ***, P < .0005).

To confirm that our results were not an artifact of CRISPR screening, we repeated the above analysis using RNAi data from 497 cancer cell lines. Consistent with the results obtained using CRISPR, we found that cancer cell lines harboring deletions of chromosome 1p36 exhibited a significant increase in vulnerability to CDK11-targeting shRNAs (Fig. 4E). Finally, we performed a PRISM drug screen, in which we determined the sensitivity of 835 different cancer cell lines to treatment with OTS964^46^. Consistent with the results obtained using CRISPR and RNAi, we found that cancer cell lines harboring chromosome 1p36 deletions were significantly more sensitive to OTS964 compared to other cancer cell lines (Fig. 4F and Table S10). In total, these results provide three independent lines of evidence that chromosome 1p36 deletions are associated with enhanced sensitivity to CDK11 ablation.

To our knowledge, there are two anti-cancer therapies that have received FDA approval based on a chromosomal deletion biomarker: lenalidomide, which is used in myelodysplastic syndromes that harbor a heterozygous deletion of chromosome 5q, and venetoclax, which is used in chronic lymphocytic leukemia patients that harbor a heterozygous deletion of chromosome 17p^47,48^. We noted that the overall increase in sensitivity in the 1p36-deleted cancers treated with OTS964 was modest (Fig. 4F). However, we performed the same biomarker analysis comparing lenalidomide sensitivity with 5q copy number and venetoclax sensitivity with 17p copy number, and we found that the difference between deleted and neutral/gain cell lines was smaller than the difference that we observed with OTS964 and 1p36 (Fig. S7). We conclude that the increase in OTS964 sensitivity exhibited by cancers with 1p36 deletions is consistent with the profile exhibited by FDA-approved drug/biomarker combinations in this specific assay.

### Decreased expression of CDK11 and its activating cyclin increase sensitivity to CDK11 inhibition

We next sought to uncover the biological basis for the link between 1p36 copy number and OTS964 sensitivity. We noted that the chromosome 1p36 locus encodes *CDK11A*, *CDK11B*, and *CCNL2* (Cyclin L2), which activates CDK11 function (Fig. 4G)^8^. The copy number and expression of *CDK11A*, *CDK11B*, and *CCNL2* were individually associated with enhanced sensitivity to CDK11 targeting (Fig. 4A and 4C). We speculated that cancers harboring deletions of the 1p36 locus express low levels of CDK11 and Cyclin L2, thereby rendering them more sensitive to inhibition of the remaining protein. Consistent with this hypothesis, 1p36 deletions were associated with decreased expression of *CDK11A*, *CDK11B*, and *CCNL2* in both the Cancer Cell Line Encyclopedia and TCGA datasets (Fig. S8A-B).

We subsequently sought to explore the frequency of 1p36 deletions in human cancers. We found that these deletions were common across diverse cancer types: within the TCGA, 25% of cancers exhibited 1p36 deletions, comparable to the frequency of deletions in *PTEN* (25%) and *RB1* (31%)(Fig. S8C). We believe that CDK11 itself is not the target of these deletions. Indeed, we found that no cancers in TCGA harbor a homozygous deletion of *CDK11A*, *CDK11B*, or *CCNL2*, which is consistent with our data demonstrating that CDK11 is essential for tumor proliferation (Fig. S8D and Table S11). However, several other genes on chromosome 1p are subject to homozygous deletions, including *CDKN2C* and *PARK7*. GISTIC2.0, an algorithm designed to identify driver genes within somatic copy number alterations, confirmed that the *CDK11*/*CCNL2* locus is commonly deleted across cancer types (Q < 10^-213^), with particularly high levels in epithelial cancers, including breast and colorectal adenocarcinoma (Table S12)^49^. However, *CDK11*/*CCNL2* is not located in a deletion peak across cancer types; the nearest peak in 1p36 spans from *VAMP3* to *RERE*. Finally, manual inspection of the *CDK11/CCNL2* locus revealed several additional tumor suppressors that were located nearby, including *TP73*, *CHD5*, and *RPL22* (Fig. 4G). We speculate that deletions of many of these genes may be under positive selection during tumor evolution, which can as a consequence result in collateral heterozygous deletions of *CDK11/CCNL2*.

To establish a causative relationship between 1p36 copy number and sensitivity to CDK11 inhibition, we used CRISPR to generate heterozygous segmental deletions of the 1p36 locus in the near-diploid cancer cell lines, A2780 and DLD1 (Fig. 4G). We found that deleting a single copy of 1p36 caused a significant increase in sensitivity to OTS964 in each cell line when compared to their respective parental cell lines and to clones that were transfected with the same gRNAs but that did not acquire a 1p36 deletion (Fig. 4H). To confirm that these effects were driven by decreased expression of CDK11 and Cyclin L2, we over-expressed *CDK11B* and *CCNL2* in wild-type and 1p36-del A2780 cells. The over-expression of these genes had no effect on OTS964 sensitivity in 1p36-neutral cells but restored OTS964 sensitivity to wild-type levels in the 1p36-del cell line (Fig. 4I). We conclude that deletions of the 1p36 locus enhance sensitivity to both genetic and pharmacological ablation of CDK11 by decreasing the copy number of the *CDK11* and *CCNL2* genes.

### CRISPR screening verifies that targeting CDK11 and Cyclin L enhance sensitivity to CDK11 inhibition

As an independent approach to uncover biomarkers conferring increased sensitivity to CDK11 inhibition, we performed a chemogenetic interaction screen using OTS964. We transduced the near-diploid leukemia cell line NALM6 with a genome-wide CRISPR library and then cultured the cells in normal media or in media containing a sub-lethal dose of OTS964 for eight days (Fig. S9A). Under these conditions, the cells grown in normal media underwent 7.5 population doublings while the cells cultured in OTS964 underwent 6.4 population doublings. We compared gRNA dropout between the OTS964-treated and untreated cells, and we identified the loss of *CCNL1* (Cyclin L1) as the alteration causing the strongest increase in OTS964 sensitivity (Fig. S9B-C and Table S13A). Additionally, *CDK11A* and *CDK11B* were both among the top 70 genes whose loss enhanced the effects of OTS964 treatment. Gene Set Enrichment Analysis revealed that genetic ablation of splicing-associated factors was also significantly associated with increased sensitivity to OTS964, consistent with CDK11’s role in this pathway (Fig. 2, S9D, and Table S13B-C). In total, these results provide independent evidence that sensitivity to CDK11 inhibition is enhanced by the loss of the genes that encode CDK11 and/or Cyclin L.

### Increased sensitivity to CDK11 inhibition in patient-derived breast cancer organoids with 1p36 deletions

Finally, we sought to determine whether the association between 1p36 copy number and sensitivity to CDK11 inhibition extended to primary human cancer specimens. Toward that goal, we assessed a panel of primary patient-derived breast cancer resections that were minimally-passaged and cultured as 3-D organoids (Fig. S10A)^50^. We determined the copy number of the 1p36 locus using low-pass whole-genome sequencing, and we identified four organoids that harbored a heterozygous deletion of this locus and eight organoids in which this locus was copy-neutral or amplified. Next, we calculated IC50 values for OTS964 in each organoid, and we found that the organoids harboring a 1p36 deletion were 2.5-fold more sensitive to OTS964 compared to the organoids that were neutral or amplified at this locus (36 nM vs. 88 nM, P < .04; two-tailed t-test)(Fig. S10B-C). These results provide additional data demonstrating a link between 1p36 copy number and OTS964 sensitivity in primary patient-derived cancer specimens.

### Characterization of MEL-495R, a CDK11 inhibitor suitable for *in vivo* dosing

We sought to determine the safety and efficacy of CDK11 inhibition as a therapeutic strategy. However, an initial ADME profile of OTS964 revealed several liabilities that could hinder its successful use *in vivo* (Fig. S11A). We therefore tested various derivatives of OTS964 to identify potential CDK11 inhibitors with improved metabolic properties. We developed MEL-495, which exhibited a significant increase in solubility and half-life and a decrease in plasma protein binding relative to OTS964 (Fig. S11 and S12). Next, we isolated the pure *R* enantiomer of MEL-495 and compared this compound (MEL-495R) with racemic MEL-495 (Fig. S13). We found that the IC50 of MEL-495R was comparable to the IC50 of OTS964 and was approximately two-fold lower than the IC50 of MEL-495, indicating that this enantiomer is largely responsible for the compound’s activity against CDK11 (Fig. S14A). We therefore used the pure MEL-495R enantiomer in our subsequent experiments.

Next, we performed a KINOMEscan assay to measure the binding between MEL-495R and 468 human kinases. We found that 1 µM MEL-495R resulted in 99.6% engagement of CDK11B and 95.3% engagement of CDK11A (Fig. S14B and Table S14). MEL-495R exhibited an s(10) selectivity score of 0.051, indicating strong selectivity for CDK11. To verify that MEL-495R targets CDK11 *in cellulo*, we determined its IC50 values in a melanoma cell line (A375) and a breast cancer cell line (MDA-MB-231) that expressed either wild-type CDK11B or CDK11B^G579S^. We found that MEL-495R exhibited IC50 values in the 20-25 nM range in the wild-type cell lines while the expression of CDK11B^G579S^ caused a ∼5-fold increase in MEL-495R resistance (Fig. S14C). Finally, we assessed the *in vivo* pharmacokinetics of MEL-495R, which revealed favorable drug-like properties (Fig. S14D). MEL-495R demonstrated good oral bioavailability (21.85%) with rapid absorption (T_max_ = 0.5h) and moderate plasma half-life (t_1/2_ = 2-3h). The volume of distribution (12.86 L/kg) suggested extensive tissue distribution. These properties supported the further evaluation of MEL-495R as an orally available CDK11 inhibitor.

### Generation of a mouse model to differentiate between on-target and off-target toxicity of CDK11 inhibitors

To characterize CDK11 as a possible target for cancer treatment, we sought to determine whether CDK11 inhibition results in a tolerable safety profile *in vivo*. However, investigating the toxicity of a novel drug target represents a major challenge in molecular pharmacology, as distinguishing between “on-target” and “off-target” toxicity is extremely challenging. While our prior experiments revealed that MEL-495R exhibits strong selectivity for CDK11 *in vitro*, these findings are insufficient to conclude that any toxicity resulting from MEL-495R administration *in vivo* is necessarily a consequence of CDK11 inhibition. Firstly, our KINOMEscan analysis revealed that MEL-495R interacts with several other kinases that could impact organismal physiology, including CDK7, CIT, and GSK3B (Fig. S14B and Table S14). Secondly, many kinase inhibitors cause toxicity by inhibiting proteins outside of the kinase family, and it is not biologically possible to individually assess the interaction between a small-molecule and the thousands of non-kinase proteins encoded in a mammalian genome^51^. Thirdly, the *in vivo*-specific metabolism of a drug can produce derivative compounds that exhibit unique pharmacological interactions and can influence tolerability. To address this uncertainty in interpreting *in vivo* experiments with MEL-495R, we sought to develop an approach to determine whether any toxicity resulting from treatment with this drug was a direct consequence of CDK11 inhibition.

We hypothesized that we could establish a genetically-encoded CDK11 inhibitor resistance model to differentiate between on-target and off-target drug toxicity *in vivo*. Toward that goal, we identified a mutation in mouse Cdk11b, Cdk11b^G568S^, that is orthologous to the human mutation that we previously characterized (Fig. 5A). We used CRISPR-mediated homology-directed repair to knock this mutation into mouse zygotes and achieved germline transmission of the mutant allele (Fig. 5B). We found that mice with the Cdk11b^G568S/WT^ genotype were born at the expected Mendelian ratio and gained weight at wild-type rates, indicating that this mutation is broadly tolerated (Fig. 5C-D). Next, we isolated mouse embryonic fibroblasts (MEFs) with Cdk11b^WT/WT^ and Cdk11b^G568S/WT^ genotypes. We found that the expression of Cdk11b^G568S^ caused a 9-fold decrease in MEF sensitivity to MEL-495R, verifying that this mutation blocks the ability of MEL-495R to inhibit mouse CDK11 function (Fig. 5E).

**Figure 5.**
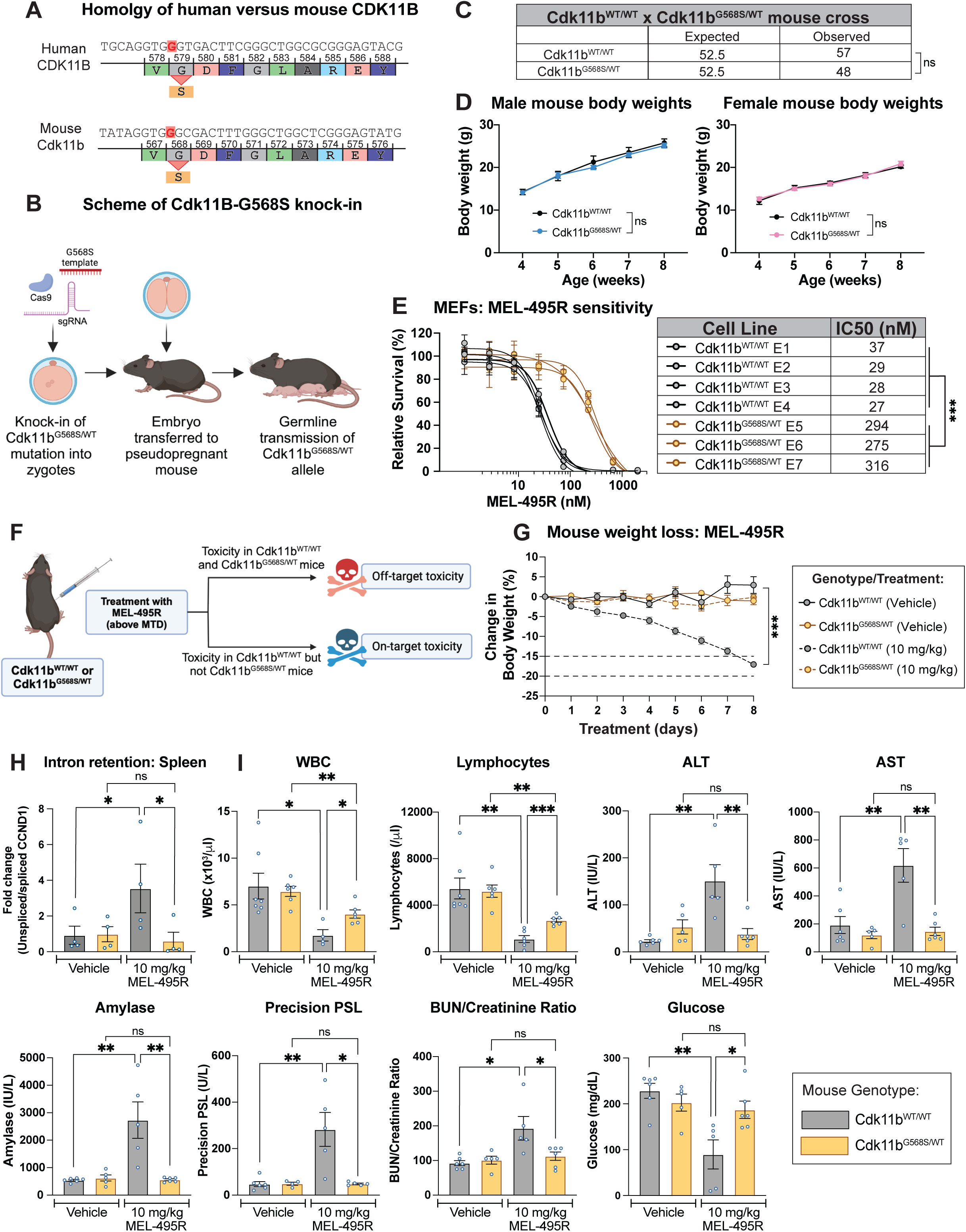
Derivation and analysis of a genetically-engineered mouse model for differentiating between on-target and off-target toxicity for CDK11 inhibitors. A) Schematic of the homology between human and mouse CDK11 around the location of the G579S resistance mutation. B) Diagram of the strategy used to knock-in the Cdk11b^G568S^ mutation into mouse zygotes. C) Mendelian ratio of genotypes of the progeny from crosses between Cdk11b^WT/WT^ and Cdk11b^G568S/WT^ mice. Cdk11b^WT/WT^ and Cdk11b^G568S/WT^ progeny were recovered at the expected frequencies. Statistical significance was determined by Fisher’s exact test (*, P < .05; **, P < .005; ***, P < .0005). D) Body weights of male and female Cdk11b^WT/WT^ and Cdk11b^G568S/WT^ mice. Mean ± SEM from 4 to 9 mice per group. Statistical significance was determined by unpaired t-tests (*, P < .05; **, P < .005; ***, P < .0005). E) Cdk11b^WT/WT^ and Cdk11b^G568S/WT^ MEFs were treated with varying doses of MEL-495R (left panel). The IC50 values are provided in the associated table (right panel). Mean ± SEM, data from three independent replicates. Statistical significance was determined by unpaired t-tests (***, P < .0005). F) Schematic of the approach to use mice expressing Cdk11b-G568S to assess the CDK11-dependent and CDK11-independent consequences of MEL-495R treatment. G) Female Cdk11b^WT/WT^ and Cdk11b^G568S/WT^ mice were split into 4 groups, treated with vehicle or 10 mg/kg MEL-495R daily, and body weights were recorded. Mean ± SEM of data from 4 mice per group. Statistical significance was determined by unpaired t-tests (***, P < .0005). H) Bar graph displaying fold change of unspliced/spliced CCND1 in RNA extracted from mice treated as described in F. Relative gene expression levels were normalized to B2M as a reference gene. Mean ± SEM, data from three independent replicates. Statistical significance was determined by unpaired t-tests (*, P < .05; **, P < .005; ***, P < .0005). I) Blood was harvested from mice upon the endpoint of the study described in (G). Complete blood counts and blood chemistry were performed from mice treated as shown in (H). Complete results are included in Table S15. Mean ± SEM of data from 4-6 mice per group. Statistical significance was determined by unpaired t-tests (*, P < .05; **, P < .005; ***, P < .0005).

### CDK11 inhibition results in substantial on-target toxicity *in vivo*

We hypothesized that we could leverage our Cdk11b^G568S^ mouse model to differentiate between on-target and off-target toxicity of CDK11 inhibitors. In particular, if MEL-495R induces toxicity in both Cdk11b^WT/WT^ and Cdk11b^G568S/WT^ mice, then this would suggest that these toxic effects are independent of CDK11 inhibition. In contrast, if the Cdk11b^G568S/WT^ mice are resistant to certain side effects compared to the Cdk11b^WT/WT^ mice, then this would indicate that those effects of MEL-495R administration are a result of CDK11 inhibition (Fig. 5F). This mouse model could therefore allow us to uncover the effects of systemic CDK11 inhibition *in vivo* and would identify any off-target effects of MEL-495R treatment.

Before using this model, we first conducted a maximum-tolerated dose (MTD) study on MEL-495R in wild-type mice. We determined that a dose of 2 mg/kg MEL-495R administered once daily (Q.D.) via intraperitoneal injection (IP) was tolerated, while higher doses resulted in weight loss and mouse morbidity (Fig. S14E). Next, we bred a large cohort of female, 8-week-old Cdk11b^WT/WT^ and Cdk11b^G568S/WT^ mice and treated them with either vehicle or with a toxic dose of MEL-495R (10 mg/kg). As expected, all of the wild-type mice receiving 10 mg/kg MEL-495R rapidly lost weight and had to be euthanized after five days of treatment, while wild-type and Cdk11b^G568S/WT^ mice tolerated the vehicle injections without significant weight loss. Remarkably, the Cdk11b^G568S/WT^ mice treated with MEL-495R did not lose any weight relative to the vehicle-treated mice and did not exhibit any clinical symptoms requiring euthanasia (Fig. 5G). These results establish that the severe toxicity resulting from MEL-495R treatment is broadly an on-target, CDK11-driven effect.

We euthanized the mice from this experiment and comprehensively evaluated the effects of MEL-495R. As an *in vivo* biomarker to assess CDK11 inhibition, we isolated RNA from mouse spleens and quantified intron retention via qPCR. We found that MEL-495R treatment resulted in a significant increase in intron retention in Cdk11b^WT/WT^ but not Cdk11b^G568S/WT^ mice, confirming that the expression of the G568S allele preserved CDK11 function in animals treated with MEL-495R (Fig. 5H). Next, we performed comprehensive hematological and clinical chemistry analysis on the mice. We found evidence of significant multi-organ system dysregulation caused by MEL-495R treatment. The compound induced severe immunosuppression, characterized by a 77% reduction in total white blood cells, primarily driven by an 85% decrease in lymphocytes and an 82% reduction in eosinophils (Fig. 5I, S15, and Table S15). MEL-495R treatment also resulted in marked hepatocellular injury, evidenced by substantial elevations in the liver enzymes AST and ALT. Additionally, the compound induced significant metabolic perturbations, including hypoglycemia, hypercholesterolemia, and an elevated BUN/creatinine ratio, suggesting impacts on glucose homeostasis and kidney function. Nearly all of these abnormalities were rescued by the expression of the Cdk11b resistance mutation: Cdk11b^G568S/WT^ mice that were treated with MEL-495R exhibited no significant abnormalities in AST, ALT, amylase, cholesterol, or BUN/creatinine. MEL-495R treatment still resulted in leukopenia in Cdk11b^G568S/WT^ mice, though not as severe as observed in wild-type mice. Hematopoietic toxicity has previously been documented after treatment with other CDK inhibitors, and we speculate that some of this leukopenia may result from residual activity of MEL-495R against other members of the CDK family^52–55^. In total, our analysis demonstrates that MEL-495R causes broad-spectrum organ dysfunction, and this pathology is a direct consequence of inhibiting CDK11.

### Tolerable doses of MEL-495R do not result in substantial anti-cancer activity *in vivo*

Many cancer drugs cause severe side effects, but they are still able to induce tumor regressions at doses below their toxicity thresholds. We conducted a series of mouse xenograft and allograft experiments to determine whether tolerable doses of MEL-495R could slow or prevent tumor growth (Fig. 6A). We initially utilized a dose of 2 mg/kg MEL-495R, which is 5-fold lower than the toxic dose that was tested in Figure 5 and is the maximum dosage that did not result in significant weight loss in wild-type mice (Fig. S14E). We also tested higher doses delivered every third day (5 mg/kg) or delivered orally (45 mg/kg). We selected cell lines from diverse cancer types that each exhibited an MEL-495R IC50 value below 25 nM, including two cell lines with 1p36 deletions (Fig. 6A). We used nude mice as a host organism for all experiments involving human cancer cell lines, and we used C57BL/6 as a host organism for the B16F10 mouse melanoma cell line.

**Figure 6.**
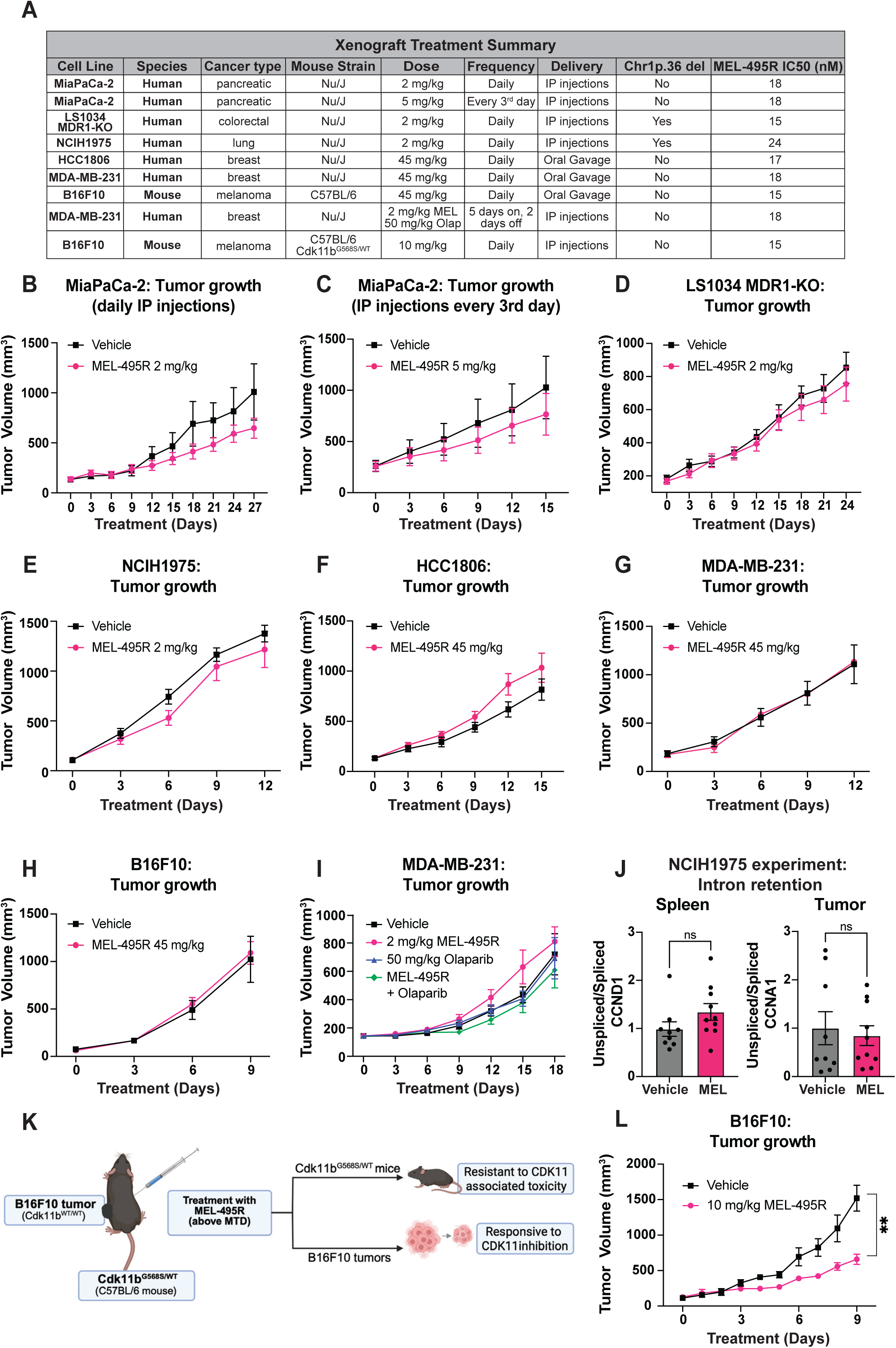
Tolerable doses of MEL-495R exhibit minimal efficacy against human and mouse cancer xenografts. A) A table summarizing the xenograft experiments performed with MEL-495R. B-I) Tumor growth measurements for the xenograft experiments described in A. J) As a biomarker for CDK11 inhibition, we assessed intron retention in the spleen (left) and the tumor (right) from the experiment in (E). K) A schematic of the experiment to determine whether on-target toxicity limits the anti-cancer efficacy of MEL-495R. Mouse B16F10 cells were injected into Cdk11b^G568S/WT^ mice and allowed to form xenografts. The mice were subsequently treated with a dose of MEL-495R that is toxic in wild-type mice but well-tolerated in Cdk11b^G568S/WT^ mice. L) Tumor growth measurements from the xenograft experiment described in K. Statistical significance was determined by unpaired t-test (**, P < .005).

Across these diverse conditions, we found that tolerable doses of MEL-495R resulted in minimal or no suppression of tumor growth (Fig. 6B-I). For instance, we injected mice with the human lung cancer cell line NCIH1975, which harbors a deletion in 1p36 and exhibits an MEL-495R IC50 value of 24 nM *in vitro*. However, after 12 days of treatment, we found that vehicle-treated mice harbored an average tumor size of 1,376 ± 85 mm^3^ while MEL-495R-treated mice harbored an average tumor size of 1,216 ± 181 mm^3^ (Fig. 6E). We obtained similar results when treating mice with xenografts derived from pancreas, breast, colon, and melanoma cell lines.

We previously observed that CDK11 inhibition exhibited synergistic cell killing of the triple-negative breast cancer cell line MDA-MB-231 with a PARP inhibitor *in vitro* (Fig. S5). We therefore sought to investigate whether co-administration of MEL-495R and olaparib could produce regressions of MDA-MB-231 xenografts *in vivo*. We treated mice with either vehicle, 2 mg/kg MEL-495R, 50 mg/kg olaparib, or 2 mg/kg MEL-495R + 50 mg/kg olaparib, dosed for five consecutive days followed by a two-day break. However, we did not observe a significant difference in tumor size between mice treated with vehicle and mice treated with any of these drugs or drug combinations (Fig. 6I).

To uncover the cause(s) of the minimal *in vivo* activity that we observed with MEL-495R administration, we isolated spleens and tumors from the NCIH1975 xenograft experiment described above and assessed CDK11-dependent splicing in each sample. While our toxicity experiments revealed that treatment with 10 mg/kg MEL-495R resulted in a significant increase in intron retention in mouse spleens (Fig. 5H), we found that 2 mg/kg MEL-495R had no effect on splicing in this organ or in xenografts from treated mice (Fig. 6J). These results indicate that this dosage of MEL-495R does not result in pronounced target inhibition *in vivo*, likely explaining why these treatments did not cause either organismal toxicity or tumor regression.

Finally, we sought to directly confirm that on-target toxicity limits the *in vivo* efficacy of MEL-495R. Toward that end, we bred a large cohort of age- and sex-matched Cdk11b^G568S/WT^ mice and then injected them with B16F10 mouse melanoma cells (Fig. 6K). After tumor formation, we dosed the mice with 10 mg/kg MEL-495R, which we previously found was well-tolerated by mice that expressed Cdk11b^G568S^ (Fig. 5G). While treatment of B16F10 xenografts in Cdk11b^WT/WT^ mice with 2 mg/kg MEL-495R did not affect tumor growth (Fig. 6H), treatment of these same xenografts in Cdk11b^G568S/WT^ mice with 10 mg/kg MEL-495R resulted in a significant 57% decrease in tumor size compared to vehicle-treated mice (Fig. 6L). Thus, by uncoupling MEL-495R’s on-target toxicity from its on-target anti-cancer activity, we found that MEL-495R is capable of impacting cancer growth at doses that are otherwise intolerable.

## DISCUSSION

In this work, we resolve conflicting reports regarding CDK11’s role in cancer by definitively establishing that its kinase activity is required for cancer cell viability. CDK11 was initially characterized as a tumor suppressor due to its frequent deletion in cancer and its association with apoptosis^56–58^. However, our findings indicate that CDK11 is likely not the target of the recurrent 1p36 deletions, as the CDK11 locus is not subject to either focal or homozygous deletions. Instead, the loss of CDK11 may occur as a collateral consequence of the deletion of other tumor suppressors encoded on chromosome 1p, including *CDKN2C*, *CHD5*, and *TP73*. Interestingly, many previous large-scale analyses using CRISPR or RNAi have failed to identify CDK11 as a cancer-essential gene^44,59–62^. Here, we designed (Fig. 1A) and analyzed (Fig. S6A) gRNAs that recognize both *CDK11A* and *CDK11B*, and we showed that the expression of either paralog is sufficient for cellular viability. This redundancy between paralogs likely prevented the identification of CDK11 in prior efforts to map cancer vulnerabilities.

Our re-analysis of published CRISPR screening data, drug sensitivity profiling using OTS964, and experiments in primary breast cancer organoids consistently identified 1p36 deletions as a genomic alteration that significantly enhances sensitivity to CDK11 inhibition. We further demonstrated that the over-expression of *CDK11B* and *CCNL2* in 1p36-deleted cells restores a wild-type level of resistance to OTS964. Put together, these data identify CDK11 as a new “CYCLOPS” gene - a class of cancer drug targets for which partial loss of the genomic region encoding that target creates a unique vulnerability to inhibition of the remaining protein^63^. The CYCLOPS concept has been successfully translated to the clinic, with the widespread use of lenalidomide to target 5q-del myelodysplastic syndrome based on the decreased copy number of CK1α^64^. The prevalence of 1p36 deletions across diverse cancer types (∼25% of tumors in TCGA) suggests that a substantial patient population could exhibit enhanced sensitivity to CDK11 inhibition.

To explore the utility of CDK11 as a cancer vulnerability, we developed an orally available CDK11 inhibitor called MEL-495R and characterized its effects *in vivo*. As the therapeutic potential of CDK11 has not been previously explored, we sought to establish an approach that would allow us to rigorously differentiate between the CDK11-dependent and CDK11-independent effects of MEL-495R treatment. Notably, off-target drug toxicity has become increasingly recognized as a major source of problems in oncology drug development^26,65–69^. The mischaracterization of compound selectivity can lead to unanticipated side effects and prevent the selection of a patient population that is most likely to respond to a particular agent. Even for compounds that are clinically marketed, off-target activity against both kinases and non-kinase targets can produce toxic sequelae that limit effective dosing regimens^69–71^. A comprehensive evaluation of drug selectivity allows for the rational design of dosing strategies, improved prediction of therapeutic index, and informed selection of responsive patient populations.

Currently, most techniques for evaluating compound selectivity are performed *in vitro* or in cell culture. Differentiating between the “on-target” and “off-target” effects of a drug candidate *in vivo* represents an unsolved problem in cancer pharmacology. To overcome this limitation, we generated a mouse model based on the OTS964-resistance mutation that we previously discovered^26^. As this mutation blocks inhibitor binding to CDK11^28^, we reasoned that any remaining effects of the drug in mutant mice represent a CDK11-independent interaction. We speculate that this same approach could be used to investigate the *in vivo* selectivity and toxicity of any experimental cancer drug with a resistance-granting mutation, so long as that mutation by itself does not markedly alter mouse physiology.

Our experiments using the Cdk11b^G568S^ mouse model revealed that CDK11 inhibition resulted in a significant, multi-organ pathology. This morbidity may reflect CDK11’s conserved role in several fundamental processes, including transcription, splicing, and DNA repair. Indeed, our work elucidates a dual mode of gene expression regulation by CDK11, in which it promotes the transcription of certain genes by activating RNAPII and ensures accurate splicing of others by phosphorylating SF3B1. We discovered that the maximum tolerated dose of MEL-495R resulted in minimal impact on intron retention levels in both tumor xenografts and normal mouse tissues, indicating insufficient target engagement to drive efficacy at doses that could be safely administered. Future work on CDK11 as a therapeutic target could therefore seek to enhance tumor-specific drug accumulation through ligand-targeted drug conjugates, alternative delivery strategies, or other therapeutic modifications.

## ACKNOWLEDGMENTS

Research in the Sheltzer Lab is supported by NIH grants R01CA237652 and R01CA276666, Department of Defense grant W81XWH-20-1-068, an American Cancer Society Research Scholar Grant, a Mark Foundation Drug Discovery Award, a sponsored research agreement from Ono Pharmaceuticals, and a sponsored research agreement from Meliora Therapeutics. Research in the Krainer Lab is supported by NIH grant GM42699.

We thank Steve Corsello (Stanford University) for providing the LS1034 MDR1-KO cells. We thank the Yale Genome Editing Center for their help in generating the Cdk11b-G568S mouse. We thank Donglai Shen and Ardiana Moustaki for assistance with experiments. We thank the Yale Animal Resources Center Staff for assistance with mouse experiments. The Yale Flow Cytometry and Precision Cancer Modeling core facilities are supported in part by an NCI Cancer Center Support Grant # NIH P30 CA016359. This work was performed with assistance from the CSHL Flow Cytometry and Sequencing Technologies & Analysis Shared Resources, which are supported in part by the Cancer Center Support Grant 5P30CA045508. We thank Suresh Jain and Intonation Research Laboratories for the initial synthesis of MEL-495 and MEL-495R.

## METHODS

### BASIC CELL CULTURE TECHNIQUES

#### Cell lines and culture conditions

The identity of each cell line was confirmed by STR profiling (University of Arizona Genetics Core, Tucson, AZ). All cell lines were grown in a humidified environment at 37°C and 5% CO_2_. Sources of each cell line and their culture conditions are listed in Table S16A.

#### Production of lentivirus

Lentivirus was generated by transfecting the plasmid of interest into HEK293 cells using the calcium phosphate or PEI methods^72^. Viral supernatant was harvested between 36-48 hours post-transfection. The supernatant was filtered using a 0.45 μM syringe and used immediately for transduction or stored at -80°C.

#### Generation of CDK11B-G579S knock-in mutant cell lines

Cell lines were transiently transfected with 2 µg of a plasmid encoding Cas9, a gRNA targeting *CDK11B* exon 16, and GFP along with 100 pmol of a single-stranded donor template to introduce the desired mutation. The donor template included multiple silent mutations in addition to the desired mutation in order to prevent re-cutting after template-mediated repair. Transfections were performed with Lipofectamine 3000. 2-3 days post transfection, GFP+ transfected cells were single cell sorted into 96-well plates for cloning. Clones were expanded and screened for integration of the desired mutation via PCR and Sanger sequencing. Plasmids used in this study are listed in Table S16B and primers used in this study are listed in Table S16C.

## PLASMID CLONING METHODS

### CRISPR plasmid cloning

Guide RNAs for CRISPR experiments were designed with Benchling (www.benchling.com). Guides were cloned into the Lenti-Cas9-gRNA-GFP vector (Addgene, #124770) using a BsmBI digestion as previously described^72^. Plasmids were transformed into Stbl3 E. coli (Thermo Fisher Scientific, Cat. No. C737303) and sequence-verified to confirm the presence of the correct gRNA. CRISPR gRNA sequences are listed in Table S16D. cDNA plasmids were obtained from Vectorbuilder, Addgene, or cloned using standard techniques. cDNA plasmids are listed in Table S16B.

## GENE EXPRESSION ANALYSIS

### RNA-seq

RNA was extracted using RNeasy Mini Kit (Qiagen; Cat. No. 74106). Purified samples were submitted to Novogene for RNA-sequencing and quantification as previously described^73^.

### Splicing analysis

Differences in splicing patterns were identified using PSI-Sigma version 2.1^74^. In short, junction reads were obtained from short-read RNA-seq data using the STAR aligner^75^ and were passed into PSI-Sigma. Pair-wise comparisons were made to quantify number of intron-retention events for each cell-line: i) Treated 100nM OTS at 6-hour timepoint vs untreated at 6-hour timepoint, and ii) Untreated at 6-hour timepoint vs untreated at 0-hour timepoint. Each comparison was run using --fmode 3 to report all events followed by filtering for relevant events (p-value < .01, ΔPSI (%) > 20).

### ChIP-seq analysis

Processed ChIP-seq data were downloaded from GSE118051 and visualized with deepTools (v3.5.6)^76^ using functions including *ComputeMatrix*, *plotHeatmap*, and *plotProfile*. Coordinates of DNA repair genes in GRCh37 were retrieved from Ensembl BioMart (v2.60.1)^77^. For each gene, regions spanning 2 kb upstream of the transcription start site (TSS) and 2 kb downstream of the transcription end site (TES) were plotted. Log2 fold changes (log2FC) of differential peaks from pSer2 RNAPII ChIP–seq were obtained using the *multiBigwigSummary* function in deepTools (v3.5.6), based on log2FC of RPGC-normalized data and differential peak coordinates comparing control and CDK11 knockdown conditions from GSE118051. Gene Ontology (GO) enrichment analysis was performed using Enrichr (https://maayanlab.cloud/Enrichr/)^78^. Barplots of log2FC values and GO enrichment results were generated using Matplotlib (v3.7.3) and Seaborn (v0.13.0) in Python 3.8^79,80^.

### qRT-PCR

RNA was extracted from cells using the RNeasy Mini Kit (Qiagen; Cat. No. 74106) or RNAplus Kit (Macherey-Nagel; Cat. No. 740984.50). RNA was converted into cDNA using SuperScript IV VILO (Invitrogen, 11756050). Quantitative PCR assays were performed using SYBR Green PCR Master Mix (Thermo Fisher Scientific, A46109) using the ViiA 7 Real-Time PCR system (Life Technologies). Ct values were obtained and analyzed using the ΔΔCt method to compare gene levels to a housekeeping gene. For intron retention qPCR assays, primers were designed using Benchling (www.benchling.com) and PrimerBLAST and the primers were designed to amplify regions spanning the exon-intron or the exon-exon junction. Ct values were obtained and analyzed using the ΔΔCt method to compare intron retention levels to a housekeeping gene. The relative expression of unspliced and spliced transcripts was quantified and expressed as a ratio to assess intron retention. The primers used for qPCR are listed in Table S16C.

### Western blotting

Cells were lysed in radioimmunoprecipitation assay (RIPA) buffer (25 mM Tris, pH 7.4, 150 mM NaCl, 1% Triton X 100, 0.5% sodium deoxycholate, 0.1% sodium dodecyl sulfate, protease inhibitor cocktail, and phosphatase inhibitor cocktail). Protein concentrations from whole cell lysates were determined using RC DC™ Protein Assay (Bio-Rad, 500–0119) and equal amounts of protein from each sample were denatured and loaded onto an SDS-PAGE gel. The proteins were transferred onto a PVDF membrane using the Trans-Blot Turbo Transfer System (Bio-Rad). Following transfer, the membrane was blocked in 5% milk or BSA in either PBS-T or TBS-T for an hour and then incubated with primary antibodies at indicated dilutions overnight at 4°C. The membrane was washed with PBS-T or TBS-T for 30 minutes and then incubated with secondary antibody at the indicated concentration for an hour at room temperature. The primary and secondary antibodies used are listed in Table S16E. To visualize proteins of interest after antibody incubation, the protein was covered with Clarity Max Western ECL (Bio-Rad, 1705062) for 1-5 minutes and imaged using a ChemiDoc Imaging System (Bio-Rad, 12003153)

### Nuclear fractionation

A375 cells were seeded on 15 cm plates at a density of 1 million cells per plate. Two days after plating, cells were treated with either DMSO or 100nM OTS964 for 6 hours. After treatments, the cells were trypsinized and half of each sample was used for whole cell protein/RNA extraction, while the other half was used for nuclear and cytoplasmic fractionation using the PARIS kit (Invitrogen; Cat. No. AM1921). The Cell Disruption Buffer and Cell Fractionation Buffer of the PARIS kit were supplemented with 100U/mL of SUPERase inhibitor (Invitrogen; Cat. No. AM2694).

## PROTEOMICS

### Sample preparation for mass spectrometry

Samples for proteome and phosphoproteome analysis were prepared as previously described^81,82^. Proteomes were extracted using a buffer containing 200 mM EPPS pH 8.5, 8M urea, 0.1% SDS and protease/phosphatase inhibitors. Following lysis, 150 µg of each proteome was reduced with 5 mM TCEP. Cysteine residues were alkylated using 10 mM iodoacetamide for 20 minutes at RT in the dark. Excess iodoacetamide was quenched with 10 mM DTT. A buffer exchange was carried out using a modified SP3 protocol^83^. Briefly, ∼1500 µg of Cytiva SpeedBead Magnetic Carboxylate Modified Particles (65152105050250 and 4515210505250), mixed at a 1:1 ratio, were added to each sample. 100% ethanol was added to each sample to achieve a final ethanol concentration of at least 50%. Samples were incubated with gentle shaking for 15 minutes. Samples were washed three times with 80% ethanol. Protein was eluted from SP3 beads using 200 mM EPPS pH 8.5 containing Lys-C (Wako, 129-02541). Samples were digested overnight at room temperature with vigorous shaking. The next morning trypsin was added to each sample and further incubated for 6 hours at 37° C. Acetonitrile was added to each sample to achieve a final concentration of ∼33%. Each sample was labelled, in the presence of SP3 beads, with ∼300 µg of TMTPro reagents. Following confirmation of satisfactory labelling (>97%), excess TMT was quenched by addition of hydroxylamine to a final concentration of 0.3%. The full volume from each sample was pooled and acetonitrile was removed by vacuum centrifugation for 1 hour. The pooled sample was acidified and peptides were de-salted using a Sep-Pak 50mg tC18 cartridge (Waters). Peptides were eluted in 70% acetonitrile, 1% formic acid and dried by vacuum centrifugation.

### Phosphopeptide enrichment

A phosphopeptide enrichment was performed using a High-Select Fe-NTA Phosphopeptide Enrichment Kit (Thermo Fisher Scientific). Dried phosphopeptides were de-salted by Stage-Tip and re-dissolved in 5% formic acid/5% acetonitrile for LC-MS/MS. The flow through from the phosphopeptide enrichment was used for total proteome profiling.

### Basic pH reversed-phase separation (BPRP)

TMT labeled peptides were solubilized in 5% acetonitrile/10 mM ammonium bicarbonate, pH 8.0 and ∼300 µg of TMT labeled peptides were separated by an Agilent 300 Extend C18 column (3.5 µm particles, 4.6 mm ID and 250 mm in length). An Agilent 1260 binary pump coupled with a photodiode array (PDA) detector (Thermo Scientific) was used to separate the peptides. A 45 minute linear gradient from 10% to 40% acetonitrile in 10 mM ammonium bicarbonate pH 8.0 (flow rate of 0.6 mL/min) separated the peptide mixtures into a total of 96 fractions (36 seconds). A total of 96 fractions were consolidated into 24 samples in a checkerboard fashion and vacuum dried to completion. Each sample was desalted via Stage Tips and re-dissolved in 5% formic acid/ 5% acetonitrile for LC-MS/MS analysis.

### Liquid chromatography separation and tandem mass spectrometry (LC-MS2)

Total proteome and phosphorylation data were collected using an Orbitrap Eclipse mass spectrometer (Thermo Fisher Scientific) coupled to a Proxeon EASY-nLC 1000 LC pump (Thermo Fisher Scientific). Peptides were separated using a 90 or 120 min gradient at 500-550 nL/min on a 35 cm column (i.d. 100 μm, Accucore, 2.6 μm, 150 Å) packed in-house. A FAIMS device was enabled during data acquisition with compensation voltages set as −40, −60, and −80 V for the total proteome samples and the phosphopeptide-enriched samples. Phosphopeptide-enriched samples were subjected to a second run in which the compensation voltages were set to −45 and −65 V^84^. For total proteome analysis, MS1 data were collected in the Orbitrap (60,000 resolution; maximum injection time set to auto; AGC 4 × 105). Charge states between 2 and 6 were required for MS2 analysis, and a 90 second dynamic exclusion window was used. Cycle time was set at 1 second. MS2 scans were performed in the Orbitrap with HCD fragmentation (isolation window 0.5 Da; 50,000 resolution; NCE 36%; maximum injection time 86 ms; AGC 1 × 105). For phosphopeptide analysis, MS1 data were collected in the Orbitrap (120,000 resolution; maximum injection time set to auto; AGC 4 × 105). Charge states between 2 and 5 were required for MS2 analysis, and a 120 second dynamic exclusion window was used. Cycle time was set at 1 second. MS2 scans were performed in the Orbitrap with HCD fragmentation (isolation window 0.5 Da; 50,000 resolution; NCE 36%; maximum injection time 250 ms; AGC 1.5 × 105).

### Data analysis

Raw files were converted to mzXML, and monoisotopic peaks were re-assigned using Monocle^85^. Searches were performed using the Comet search algorithm against a human database downloaded from Uniprot in February 2014. We used a 50 ppm precursor ion tolerance, fragment ion tolerance of 0.02, and a fragment bin offset of 0.0 for MS2 scans collected in the Orbitrap. TMTpro on lysine residues and peptide N-termini (+304.2071 Da) and carbamidomethylation of cysteine residues (+57.0215 Da) were set as static modifications, while oxidation of methionine residues (+15.9949 Da) was set as a variable modification. For phosphorylated peptide analysis, +79.9663 Da was set as a variable modification on serine, threonine, and tyrosine residues.

Each run was filtered separately to 1% False Discovery Rate (FDR) on the peptide-spectrum match (PSM) level. Then proteins were filtered to the target 1% FDR level across the entire combined dataset. Phosphorylation site localization was determined using the AScorePro algorithm^86^. For reporter ion quantification, a 0.003 Da window around the theoretical m/z of each reporter ion was scanned, and the most intense m/z was used. Reporter ion intensities were adjusted to correct for isotopic impurities of the different TMTpro reagents according to manufacturer specifications. Peptides were filtered to include only those with a summed signal-to-noise (SN) ≥ 160 across all TMT channels. An extra filter of an isolation specificity (“isolation purity”) of at least 0.5 in the MS1 isolation window was applied for the phosphorylated peptide analysis. For each protein or phosphorylation site, the filtered peptide TMTpro SN values were summed to generate protein or phosphorylation site quantification values. The signal-to-noise (S/N) measurements of peptides assigned to each protein were summed (for a given protein). These values were normalized so that the sum of the signal for all proteins in each channel was equivalent thereby accounting for equal protein loading. The resulting normalization factors were used to normalize the phosphorylation sites, again to account for equal protein loading.

To identify biological processes enriched among phosphoproteins from rescued clusters, we submitted the combined gene list to Enrichr for Gene Ontology (GO) Biological Process enrichment analysis^78^. Enrichr applied Fisher’s exact test to assess statistical significance, and the resulting enrichment terms, adjusted p-values, and odds ratios were saved in a single summary file. GO term names were cleaned by removing GO IDs for improved readability. A curated subset of terms was selected for visualization, including both high-ranking entries based on adjusted p-value and additional terms of biological interest (e.g., “Mitotic Spindle Checkpoint Signaling” and “mRNA Splicing, via Spliceosome”). Enrichment results were visualized using a horizontal bar plot, where GO terms were ordered by statistical significance.

### ANALYSIS OF OTS964 and MEL-495R SELECTIVITY

The source(s) of drugs used in this study are listed in Table S16F. The NanoBRET assay to quantify the interaction between CDK11A or CDK11B and OTS964 or palbociclib was performed as described in Wells et al.^24^ The KiNativ profiling assay was performed by ActivX Biosciences using 100 nM OTS964 and lysate from A375 cells as described in Patricelli et al.^87^ *In vitro* assays to test CDK inhibition were performed by Reaction Biology. KINOMEscan assays were performed by Eurofins.

## ANALYSIS OF CDK11 ESSENTIALITY

### CRISPR competition assays

CRISPR competition assays were performed as previously described^35^. In short, cells were transduced with a Cas9 lentivirus (Addgene #108100) and then selected with puromycin. Stable Cas9-expressing cancer cell lines were then transduced with guide RNA lentivirus supplemented with 4 μg/mL of polybrene (Santa Cruz, SC-134220). After 24 hours, the media on the cells was replenished with fresh media. Three days post-transduction, cells were passaged and the fraction of GFP+ cells was determined using a MACSQuant VYB (Miltenyi Biotec). Cells were similarly passaged and scored every three or four days, up to the 5^th^ passage, and the fold change in GFP+ cells was quantified at each passage.

### Crystal violet staining

Cells were transduced with the indicated cDNA supplemented with 4 μg/mL of polybrene (Santa Cruz, Cat. No. SC-134220). Three days post-transduction, cells were selected with 5 μg/mL blasticidin (Invivogen, ant-bl-1) or 400 μg/mL G418 (InvivoGen, Cat. No. ant-gn-1). Following selection, cells were plated in 24 well plates with the indicated concentration of OTS964. After sufficient growth, the cells were washed twice with cold PBS. The cells were then fixed for 10-30 minutes with ice-cold methanol. Cells were covered with a 0.05% crystal violet solution and incubated for 10 minutes. The crystal violet solution was then aspirated and cells were washed with PBS until excess crystal violet solution was removed. The plates were scanned and the images were cropped and contrast-adjusted in Photoshop.

## EFFECTS OF OTS964 ON DNA REPAIR

### 53BP1 foci assay

A375 cells were transduced with a construct to express a 53BP1-AppleFP fusion (Addgene, 69531). Approximately 50,000 cells were seeded into each well of 12-well plates. After 24 hours the media was replenished with fresh media containing either DMSO or the indicated dose of OTS964. After 24 hours of OTS964 pre-treatment, the media was again replaced with fresh media containing the DMSO, OTS964, doxorubicin, or OTS964 and doxorubicin. Approximately 8 hours after drug administration, the media was aspirated from the wells, washed with PBS, and the cells were fixed with 4% paraformaldehyde (PFA) (Fisher Scientific, AC416785000) in PBS pH 7.4 at room temperature for 10 minutes. Fixed cells were then washed in PBS three times followed by staining with 1ug/mL Hoechst 33342 (Invitrogen, H3570). Cells were imaged on a Cytation5 (BioTek) using 20x magnification. Foci quantification was performed manually.

### DR/NHEJ assay

U2OS EJ-DR cells^40^ were split to 6-well plates at a density of 50,000 cells per well, with triplicate wells for each treatment group. 24 hours after plating, the media was replaced with media containing the respective treatment or DMSO for a 48 hour pre-treatment period. After pre-treatment, the media was replaced with media containing the respective treatments along with 1 µM Shield-1 (Aobious, AOB1848) and 200 nM Triamcinolone (Selleck Chemicals, S1933). The mock treatment group did not receive treatment with Shield-1 and Triamcinolone. 72 hours later, the media was aspirated from wells, the cells were washed twice with PBS to remove any residual Shield-1 and Triamcinolone, and then fresh media was added. 72 hours after replacing the media, the cells were analyzed for fluorescence on a MACSQuant Flow Cytometer (Miltenyi Biotec). ΔGFP+/RFP+ was calculated by subtracting the average % GFP+/RFP+ of the mock group from each treatment group. Relative Repair Efficiency was calculated by normalizing each treatment’s ΔGFP+/RFP+ to the average ΔGFP+/RFP+ of the treatment group that had received only Shield-1 and Triamcinolone.

### PARP and CDK11 synergy assays

Cells were seeded into 6 well plates with 50,000-100,000 cells per well. 24 hours after plating, the media was replenished with fresh media containing combinations of the indicated concentrations of Olaparib and OTS964. Each drug combination was added in duplicate. After 72 hours of treatment, relative cell survival was determined by counting the cells using a Cellometer T4 (Nexcelom). Synergy calculations were performed using relative cell survival data with SynergyFinder (https://synergyfinder.fimm.fi) to obtain Bliss Synergy scores^42^. A synergy score larger than 10 is indicative of a synergistic effect.

## IDENTIFICATION OF 1P36 DELETION AS BIOMARKER

To investigate the association between 1p36 deletion and sensitivity to OTS964, we first calculated Pearson correlation coefficients between CDK11B dependency scores and both gene-level copy number and expression values across all cell lines from the DepMap database. Cell lines were then stratified by average 1p36 copy number into three groups: deep deletion (bottom 10th percentile), shallow deletion (10th–40th percentiles), and copy-neutral/gain (>40th percentile). Significant differences in mean CDK11B dependency scores across these copy number categories were assessed using Welch’s t-test (implemented via ttest_ind from the scipy library), under the assumption of unequal variances. Similar stratification and statistical testing were applied to CDK11B dependency data obtained from RNAi screens data on DepMap and OTS964-induced inhibition assays.

For gene expression-based analyses, cell lines were stratified into low expression (bottom 10th percentile), medium expression (10th–65th percentiles), and high expression (>65th percentile) groups based on CDK11B transcript levels. Differences in dependency scores across these expression categories were also evaluated using Welch’s t-test.

## 1P36 COPY NUMBER AND CDK11 INHIBITION SENSITIVITY

### 1p36 expression and copy number analysis

Analysis of 1p36 copy number gene and gene expression data was performed as described in ref^73^. In short, TCGA gene expression values were obtained from the TCGA PanCanAtlas (EBPlusPlusAdjustPANCAN_ IlluminaHiSeq_RNASeqV2.geneExp.tsv, available at https://gdc.cancer.gov/about-data/publications/pancanatlas). TCGA copy number data was obtained from the TCGA PanCanAtlas (broad.mit.edu_PANCAN_Genome_Wide_SNP_6_whitelisted.seg) and processed to link segmental copy number data with individual genes as described in refs^88,89^. Gene expression and copy number data for the Cancer Cell Line Encyclopedia was downloaded from DepMap (www.depmap.org). Homozygous deletion data was obtained from ref^90^.

### PRISM Screening

OTS964 was added to 384-well plates at 8-point dose with 3-fold dilutions in triplicate. These assay-ready plates were then seeded with 39 cell line pools of 20-25 barcoded cell lines per pool. Adherent cell pools were plated at 1250 cells per well, while suspension and mixed adherent/suspension pools were plated at 2000 cells per well in RPMI 1640 with 10% FBS (Sigma, F4135) for adherent cell pools or 20% FBS for suspension cell pools. Treated cells were incubated for 5 days then lysed. Lysate plates were collapsed together in two separate batches (cell sets) of up to 500 cell lines prior to barcode amplification and detection. As each cell line carries a unique barcode expressed as mRNA, total mRNA was captured using magnetic particles that recognize polyA sequences (Cytiva). Captured mRNA was reverse-transcribed into cDNA and then the sequence containing the unique PRISM barcode was amplified using PCR. Finally, Luminex beads that recognize the specific barcode sequences in the cell set were hybridized to the PCR products and detected using a Luminex scanner which reports signal as a median fluorescent intensity (MFI). Data processing was carried out according to code at https://github.com/cmap/dockerized_mts.

### Generation of clones harboring heterozygous deletions in chromosome 1p36

CRISPR gRNA was designed and cloned into the Lenti-Cas9-gRNA-GFP (Addgene # 124770) vector to generate heterozygous segmental deletions of the 1p36 locus (Table S16D). Cells were transfected with the targeting gRNA CRISPR plasmid using Lipofectamine 3000. After transfection, GFP+ transfected cells were single cell sorted into 96-well plates. Clones were expanded and screened for 1p36 segmental deletion via PCR and TaqMan copy number assays. Plasmids used in this study are listed in Table S16B, primers used in this study are listed in Table S16C, and TaqMan probes used in this study are listed in Table S16G.

### Cell line drug sensitivity assays

Drugs used in this study and their sources are listed in Table S16F. Cells were seeded into 96-well plates at a density of 1,000–5,000 cells per well. After 24 hours of incubation, the culture medium was replaced with fresh medium containing OTS964. Serial dilutions of the compound were prepared using 2- or 3-fold dilutions to achieve the desired concentrations. Following 72 hours of treatment, wells were washed with PBS, and fresh medium containing MTS reagent (Promega, G3582) was added. The cells were incubated with MTS at 37°C for 1–2 hours. Absorbance was measured using a plate reader at 490 nm with background correction at 666nm. Relative cell viability was determined by normalizing absorbance values to vehicle-treated control wells. IC50 values were calculated using nonlinear regression analysis in GraphPad Prism.

### Chemogenetic screen in OTS964-treated cells

The Genome-wide pooled CRISPR/Cas9 KO screen was performed by the ChemoGenix platform (IRIC, Université de Montréal; https://chemogenix.iric.ca/) as previously described^91^. Briefly, a NALM-6 clone bearing an integrated inducible Cas9 expression cassette generated by lentiviruses made from pCW-Cas9 (Addgene #50661) was transduced with the genome-wide KO EKO sgRNA library (278,754 different sgRNAs)^91^. After thawing the library from liquid nitrogen and letting it recover in 10% FBS RPMI for 1 day, KOs were induced for 7 days of culture with 2 µg/mL doxycycline. The pooled library was then split in different T-75 flasks (2.8 x 10^6^ cells per flask; a representation of 100 cells/sgRNA) in 70 mL at 4 x 10^5^ cells/mL. Cells were treated with 20 nM OTS964 for 8 days with monitoring of growth every 2 days, diluting back to 4 x 10^5^ cells/mL and adding more compound to maintain same final concentration whenever cells reached 8 x 10^5^ cells/mL. Over that period, treated cells had 6.4 population doublings whereas DMSO-only treated negative controls had 7.5. Cells were collected, genomic DNA extracted using the Gentra Puregene kit according to manufacturer’s instructions (QIAGEN), and sgRNA sequences PCR-amplified as described^91^. gRNA frequencies were obtained by next-generation sequencing (Illumina NextSeq 2000). Reads were aligned using Bowtie 2.4.4 in the forward direction only (–norc option) with otherwise default parameters and total read counts per gRNA were tabulated. Context-dependent chemogenomic interaction scores were calculated using a modified version of the RANKS algorithm^91^ that uses guides targeting similarly essential genes as controls to distinguish condition-specific chemogenomic interactions from non-specific fitness/essentiality phenotypes.

### Organoid drug sensitivity assays

OTS964 sensitivity in primary patient-derived breast cancer organoids was performed as described in Bhatia et al.^50^ and Aggarwal et al.^92^ The organoids were generated as part of the Human Cancer Models Initiative and will be made available through ATCC^93^.

## GENERATION OF THE CDK11B-G568S MOUSE MODEL

### CRISPR editing of mouse zygotes

All mouse protocols were approved by the Yale Institutional Animal Care and Use Committee (#2024-20415). Cdk11b-G568S mice were generated via CRISPR/Cas-mediated genome editing^94,95^. Potential Cas9 gRNA sequences in the vicinity of the Cdk11b-G568 codon were screened using CRISPOR^96^ and candidates were selected. Templates for gRNA synthesis were generated by PCR from a pX330 template (Addgene), then sgRNAs were transcribed *in vitro* and purified. gRNA/Cas9 RNPs were complexed and tested for activity by zygote electroporation, incubation of embryos to blastocyst stage, and genotype scoring of indel creation at the target sites. The gRNA that demonstrated the activity was selected for creating the knock-in allele (Table S16D). Guide primers for generating the template for transcription included a 5’ T7 promoter and a 3’ gRNA scaffold sequence.

A recombination template oligo (IDT) was designed to create the desired Glycine to Serine codon change; the G-to-A mutation destroys the PAM, and a silent G-to-A mutation in L572 created a new NheI restriction site for genotyping. gRNA/Cas9 RNP and the G568S template oligo were electroporated into C57Bl/6J zygotes^95^. Embryos were transferred to the oviducts of pseudopregnant CD-1 foster females using standard techniques^97^. Genotype screening of tissue biopsies from founder pups was performed by PCR amplification and Sanger sequencing to identify the desired base changes, followed by breeding and sequence confirmation to establish germline transmission of the correctly-targeted allele.

### Breeding of Cdk11b-G568S mice

To expand and maintain the Cdk11b-G568S mouse colony, backcrossing breeding cages were set up containing one male and one female. A mouse with a heterozygous Cdk11b-G568S mutation was mated with wild-type Cdk11b mice. Pups were weaned at 21 days post-birth and sexed at weaning and group-housed accordingly. Ear punches were performed to obtain tissue for genotyping and samples were sent to Transnetyx and mutational status was determined via PCR using primers as indicated in Table S16C. Routine health checks and cage changes were performed regularly. All mouse protocols were approved by the Yale Institutional Animal Care and Use Committees.

### Generation of MEFs

MEFs were isolated from embryonic 13.5 (E13.5) mouse embryos obtained from pregnant female mice. Timed pregnancies were established by mating a Cdk11b^G568S/WT^ male with a female Cdk11b^WT/WT^ C57BL/6J (Jackson Laboratories, 000664) mouse overnight, with the detection of a vaginal plug the following morning designated as embryonic day 0.5 (E0.5). Embryos were harvested from the uterus and washed in PBS. Embryonic head and organs were removed, preserving the body for fibroblast isolation. The head was retained for genotyping to confirm mutation status. The remaining tissue was minced into small fragments and further dissociated with 0.25% Trypsin-EDTA at 37°C for 10 minutes. Embryos were then further dissociated by trituration in serological pipettes of decreasing size. The cell suspension was then transferred through a 40 µm cell strainer placed on a 50 ml conical tube containing 10 ml media. Typically, MEF cultures were used before passage 5.

## ASSESSING THE TOXICITY OF CDK11 INHIBITION

### Synthesis of MEL-495 and MEL-495R

MEL-495 and MEL-495R were synthesized by Intonation Research Laboratories (Hyderabad, India). Details of the synthesis are provided in Figures S12 and S13. ADME and PK profiling were performed by Pharmaron using standard techniques. The KINOMEscan assay was performed by Eurofins as previously described^98^.

### Formulation of MEL-495R

MEL-495R was given via intraperitoneal (IP) injection or via oral gavage. For IP injections, the excipient cocktail was made up of 10% DMSO (Millipore Sigma, D2650), 40% PEG300 (Selleck, S6704), 5% Tween-80 (MedChemExpress, HY-Y1891) and 45% Saline (Quality Biological, 114-055-101). The required amount of drug was added to excipient to achieve the dose indicated. For oral gavage, the excipient cocktail was made up of 20% PEG400, 10% Cremophor, and 70% PBS, and the appropriate amount of drug was added to the excipient to achieve the dose indicated.

### Maximum tolerated dose studies

To identify the maximum tolerated dose for MEL-495R, female NU/J mice (Jackson Laboratory, 002019) aged 8-10 weeks were treated with varying concentrations of MEL-495R via IP injections. Mice were visually monitored and weighed daily to assess mouse health, and change in body weight was calculated. Mice were euthanized when 15% of initial body weight was lost or if mice displayed any signs of distress.

### Toxicity studies

To assess the toxicity of MEL-495R, female age-matched (8-12 weeks) Cdk11b^WT/WT^ and Cdk11b^G568S/WT^ mice were assigned to four experimental groups: (1) Cdk11b-wildtype vehicle-treated, (2) Cdk11b-G568S vehicle-treated, (3) Cdk11b wild-type inhibitor-treated (10 mg/kg), and (4) Cdk11b-G568S inhibitor-treated (10 mg/kg). Mice received daily treatments until they reached a humane endpoint. Throughout the study, animals were closely monitored, and body weight was recorded daily to assess treatment-related toxicity.

### Intron retention from spleen tissue

To assess intron retention in tissues following MEL-495R treatment, mice were euthanized at a humane endpoint and the spleen was harvested from mice using sterile dissection tools. 20-30 mg of tissue sample was disrupted in lysis buffer in tubes prefilled with 1.5 mm Zirconium beads (Benchmark, D1032-15). RNA from the tissues was extracted from cells using the RNeasy Mini Kit (Qiagen, 74106) or RNAplus Kit (Macherey-Nagel, 740984.50).

### Hematological/blood chemistry analysis

To assess the effects of MEL-495R treatment on hematological and serological parameters, mice were euthanized at a humane endpoint and blood was collected via cardiac puncture. Blood was collected in MiniCollect tubes with K2EDTA (Greiner, 450532) for hematology analysis or MiniCollect tubes with CAT Serum Separator (Greiner, 450533VET) for serological analysis. Hematology and serology analysis was performed by Antech Diagnostics.

## ASSESSING THE ANTI-CANCER ACTIVITY OF MEL-495R

To assess effects of MEL-495R on tumor formation, cells were harvested and resuspended at the desired concentration. For MiaPaca-2, HCC1806 and MDA-MB-231 cells, 3 million cells were injected into the right or left flank of NU/J mice (Jackson Laboratory, 002019). For LS1034-MDR1 KO cells, 4 million cells were resuspended in a 1:1 ratio with Matrigel (Corning, 356237) and injected into the left flank of NU/J mice (Jackson Laboratory, 002019). For B16F10 cells, 1 million cells were injected into the right flank of C57BL/6J mice (Jackson Laboratories, 000664). Mice were anesthetized and cells were subcutaneously injected using a 1 mL 25G x 5/8 syringe (BD, 309626). Mice were visually monitored for tumor formation routinely following injection. Once a tumor was visible, it was measured every three days by calipers. Tumor volume was calculated using the formula V = ½ x (length) x (width)^2^. Once tumor volume reached ∼100 mm^3^, treatment with MEL-495R was initiated. Tumor volume and body weight was monitored throughout the treatment period. All mouse protocols were approved by the Yale Institutional Animal Care and Use Committees.

## CODE

The code to run this analysis is available at https://github.com/sheltzer-lab/CDK11.

## SUPPLEMENTAL FIGURE LEGENDS

**Supplemental Figure 1.**
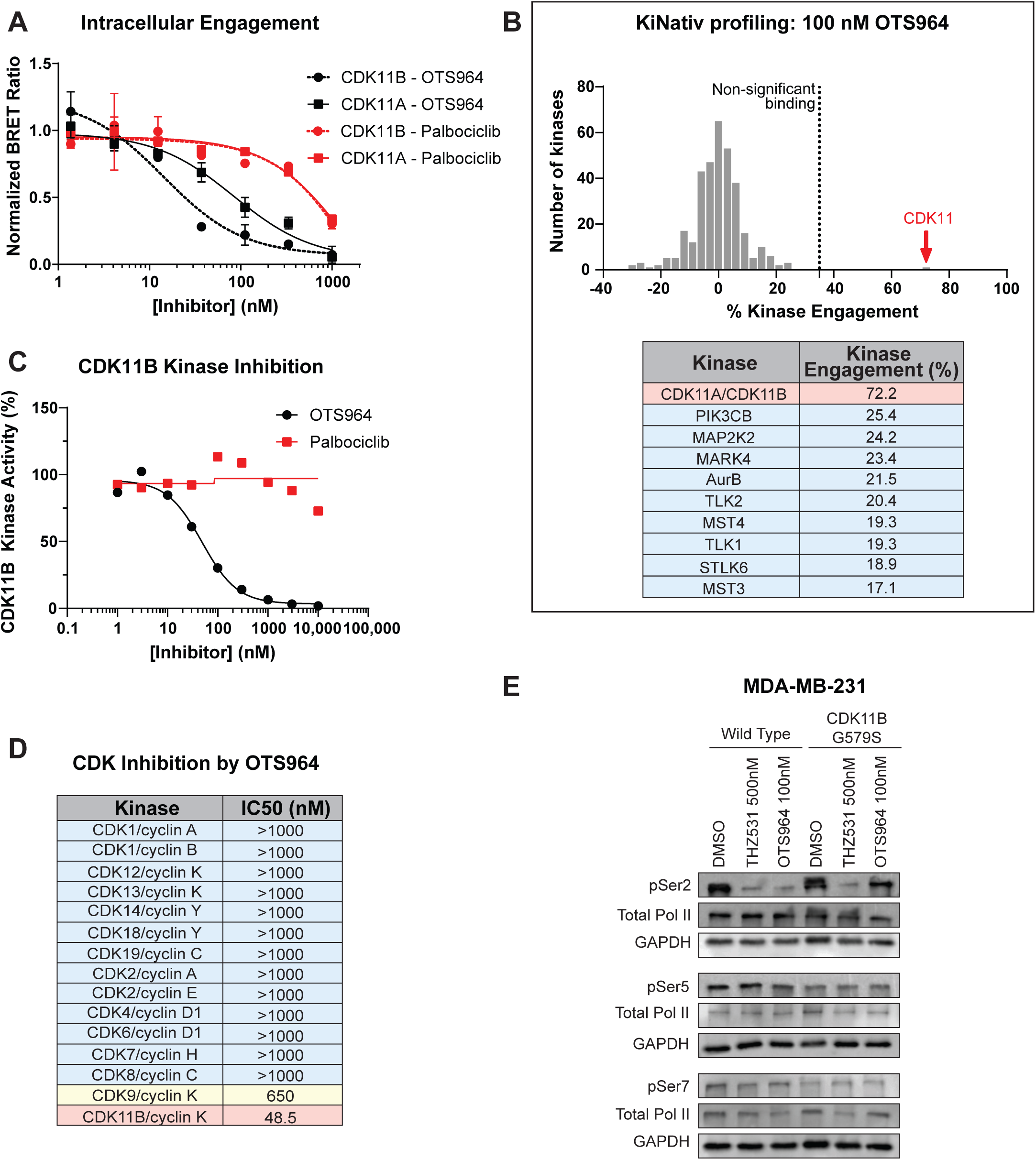
OTS964 is a selective inhibitor of CDK11 *in vitro* and *in cellulo*. A) A competitive ligand binding assay demonstrates that OTS964 binds to CDK11A and CDK11B in living cells^24^. As a control, the CDK4/6 inhibitor palbociclib was also tested and was found to exhibit substantially lower affinity for CDK11A and CDK11B. B) The top panel shows a bar graph of a KiNativ competitive ligand binding assay in which lysate from A375 cells was treated with 100 nM of OTS964 and then interactions between OTS964 and the expressed kinome were quantified. The dotted line at 35% indicates the threshold for significant binding. CDK11 is highlighted with the red arrow. The bottom panel shows a table of the top 10 hits from the KiNativ profiling and the percentage of kinase engagement. Blue indicates non-significant binding (below 35%) and red indicates significant binding (above 35%). Complete results are included in Table S1. C) CDK11B *in vitro* kinase activity assay. Dose response curves display CDK11B activity in the presence of either OTS964 or the CDK4/6 inhibitor palbociclib. D) Table summarizing OTS964’s kinase inhibition IC50 values for other CDKs as determined by Reaction Biology kinase assays. Blue indicates high IC50 values above 1 µM, yellow indicates IC50 values between 100 nM and 1 µM, and red indicates IC50 values below 100 nM. E) Western blot assessing phosphorylation status of RNAPII’s C-terminal domain at Ser2, Ser5, and Ser7 after treatment with DMSO, 500 nM of the CDK12/13 inhibitor THZ531, or 100nM OTS964. The Ser2 residue is phosphorylated by both CDK11 and CDK12/13. Expression of the G579S resistance mutation rescues the decrease in phosphorylation caused by OTS964 but not by THZ531. GAPDH levels were examined as a loading control.

**Supplemental Figure 2.**
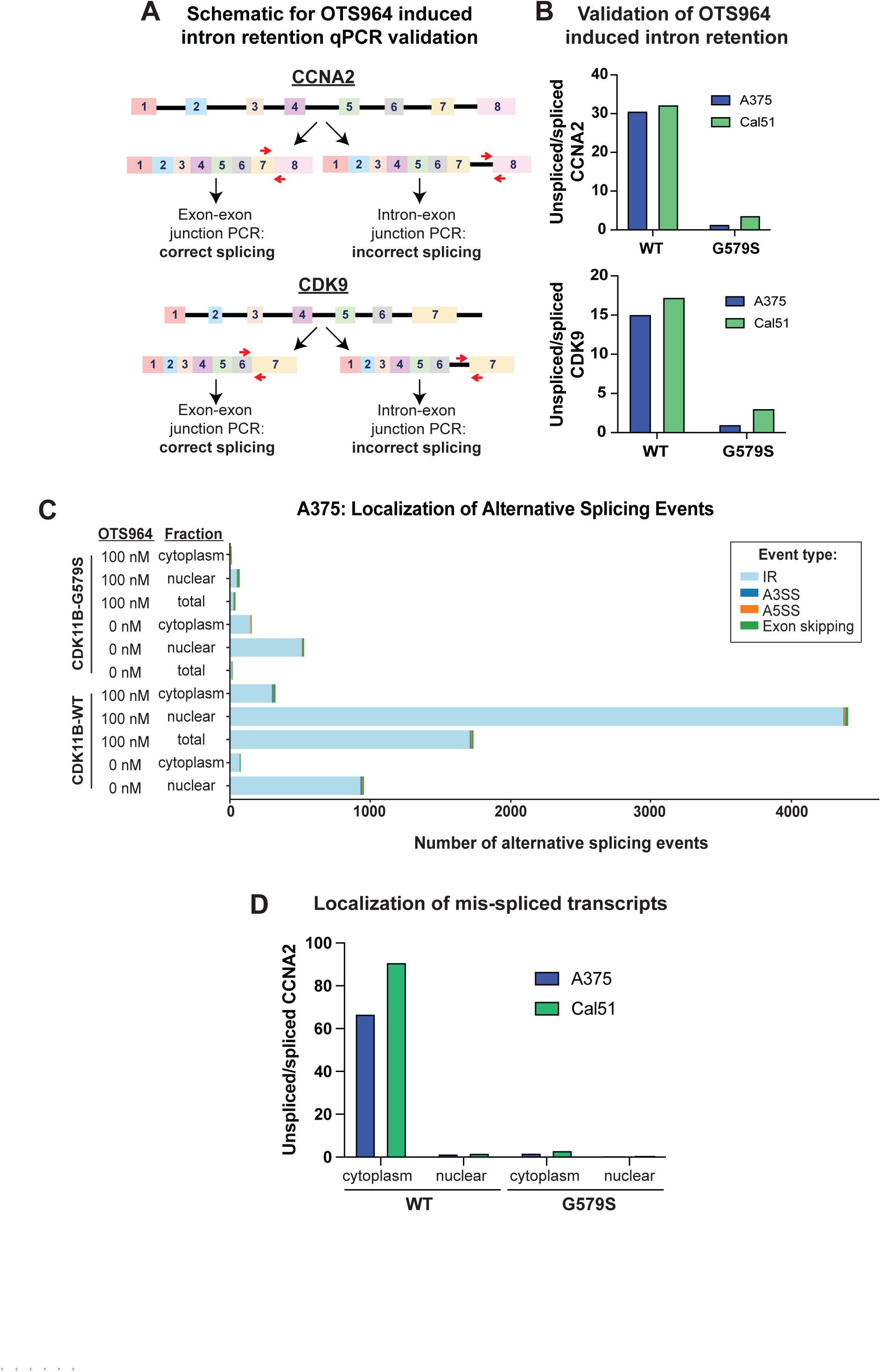
Retained introns are detected in the cytoplasm of OTS964-treated cells. A) Schematic showing the RT-qPCR method used to validate intron retention in cell lines. Primer location is indicated by the red arrows. B) Bar graph displaying fold change of unspliced/spliced *CDK9* and *CCNA2* in A375 and Cal51 cells treated with OTS964 for 6 hours. Relative gene expression levels were normalized to B2M as a reference gene. C) Bar graph showing alternate splicing events in the nuclear and cytoplasmic fractions of A375 cells expressing wild-type CDK11B or CDK11B^G579S^ treated with 100 nM OTS964 for 6 hours. Alternative splicing events were detected using PSI-Sigma with a delta-psi cutoff of 20 and p-value cutoff of 0.01^74^. IR: intron retention; A3SS: Alternative 3’ splice site; A5SS: Alternative 5’ splice site. D) Bar graph displaying fold change of unspliced/spliced *CCNA2* in nuclear and cytoplasmic fractions from A375 and Cal51 cells treated with OTS964 for 6 hours. Relative gene expression levels were normalized to B2M as a reference gene.

**Supplemental Figure 3.**
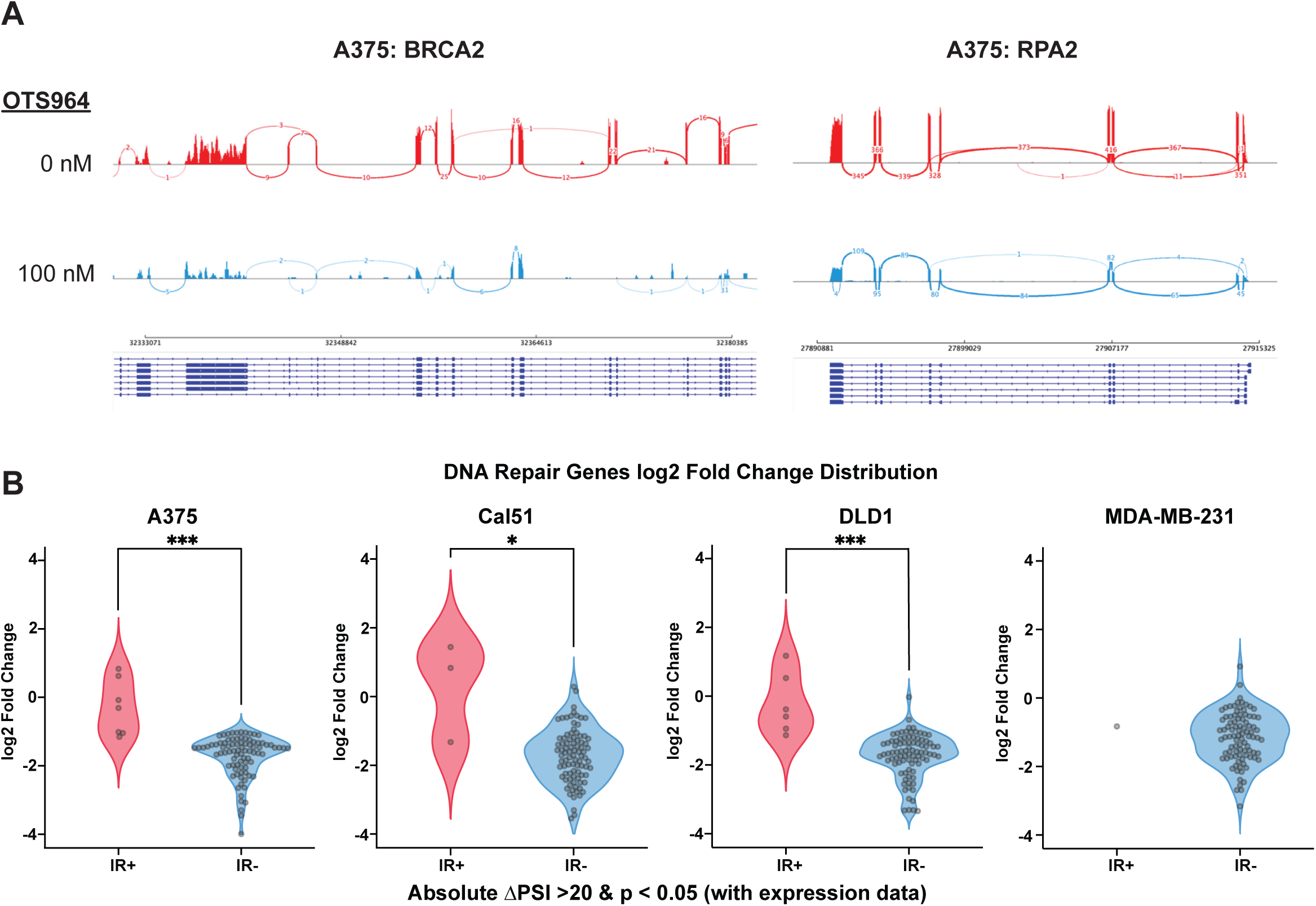
OTS964 downregulates DNA repair gene expression without inducing intron retention. A) Sashimi plots displaying the effects of OTS964 on the DNA repair genes BRCA2 and RPA2 in A375 cells. B) Violin plots displaying the expression levels of DNA repair genes that were found to exhibit intron retention (IR+) or to not exhibit intron retention (IR-) upon OTS964 treatment, as determined by PSI-Sigma. Statistical significance was determined by unpaired t-tests (*, P < .05; **, ***, P < .0005).

**Supplemental Figure 4.**
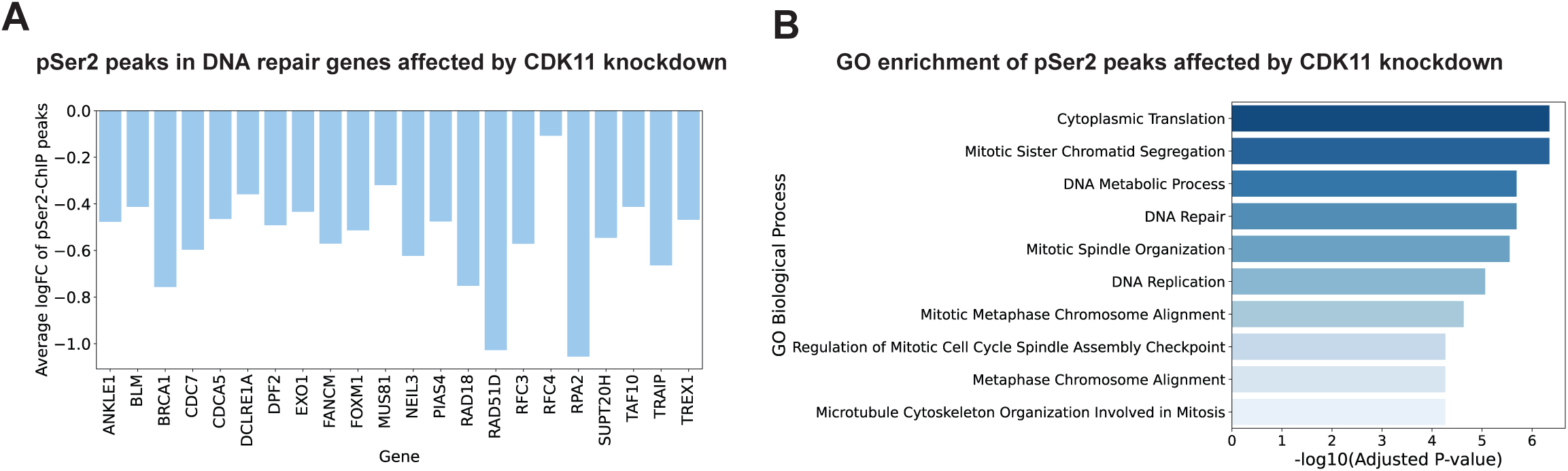
Analysis of RNAPII-pSer2 ChIP-seq peaks affected by CDK11 knockdown. A) Bar graph displaying the DNA repair genes that exhibited a significant decrease in RNAPII-pSer2 levels upon CDK11 knockdown^34^. B) Enrichment analysis of genes that exhibited a significant decrease in RNAPII-pSer2 levels upon CDK11 knockdown. Complete results are included in Table S8.

**Supplemental Figure 5.**
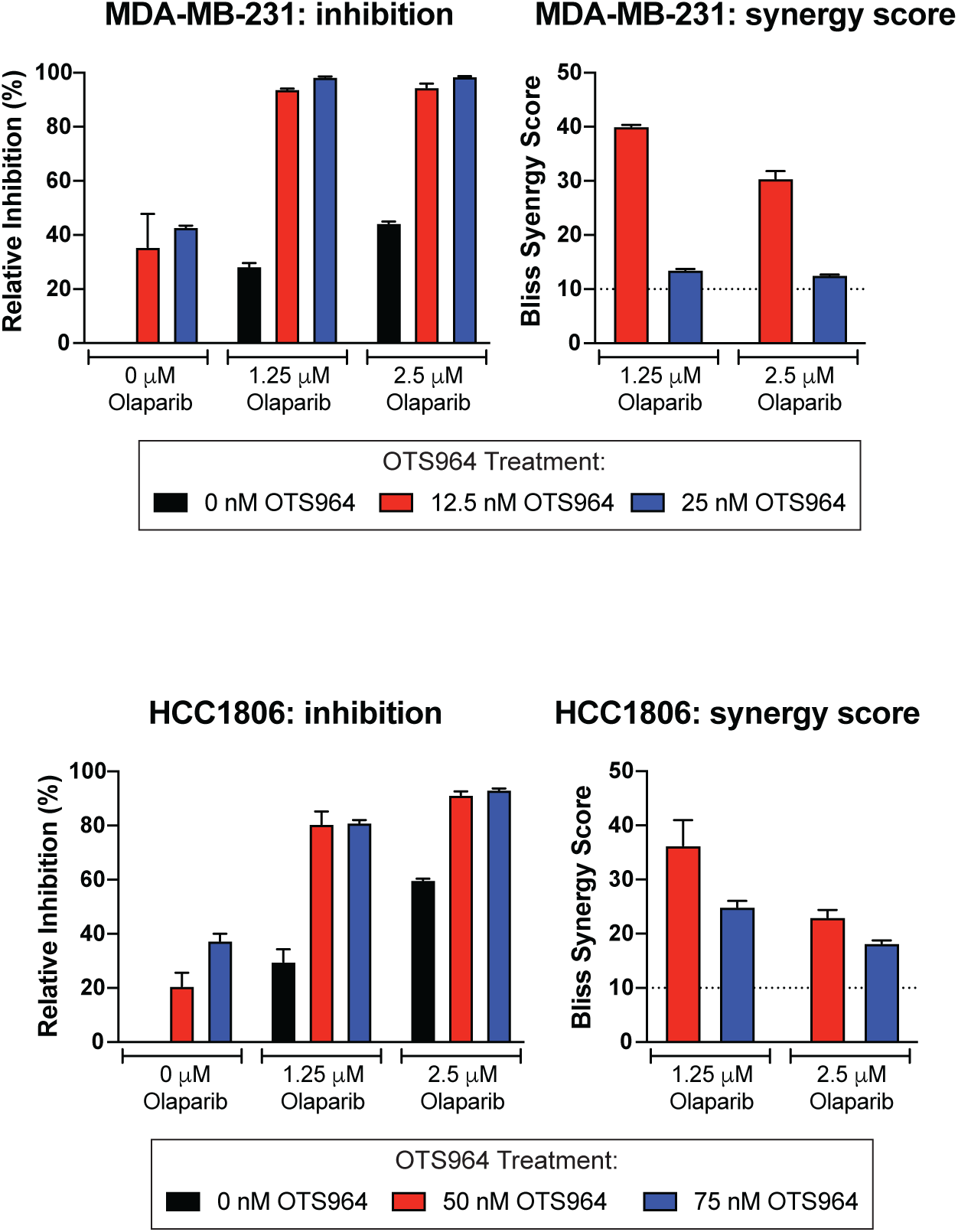
Synergistic cell killing between OTS964 and Olaparib. MDA-MB-231 and HCC1806 cells were treated with indicated doses of OTS964 and the PARP inhibitor olaparib. Relative inhibition of the cell lines was determined by normalizing treated conditions to the vehicle control and viability was assessed by manually counting the cells. Bliss synergy scores were calculated, and synergy was defined as a bliss score of >10 (dotted line). Mean ± SEM, representative data from three independent replicates.

**Supplemental Figure 6.**
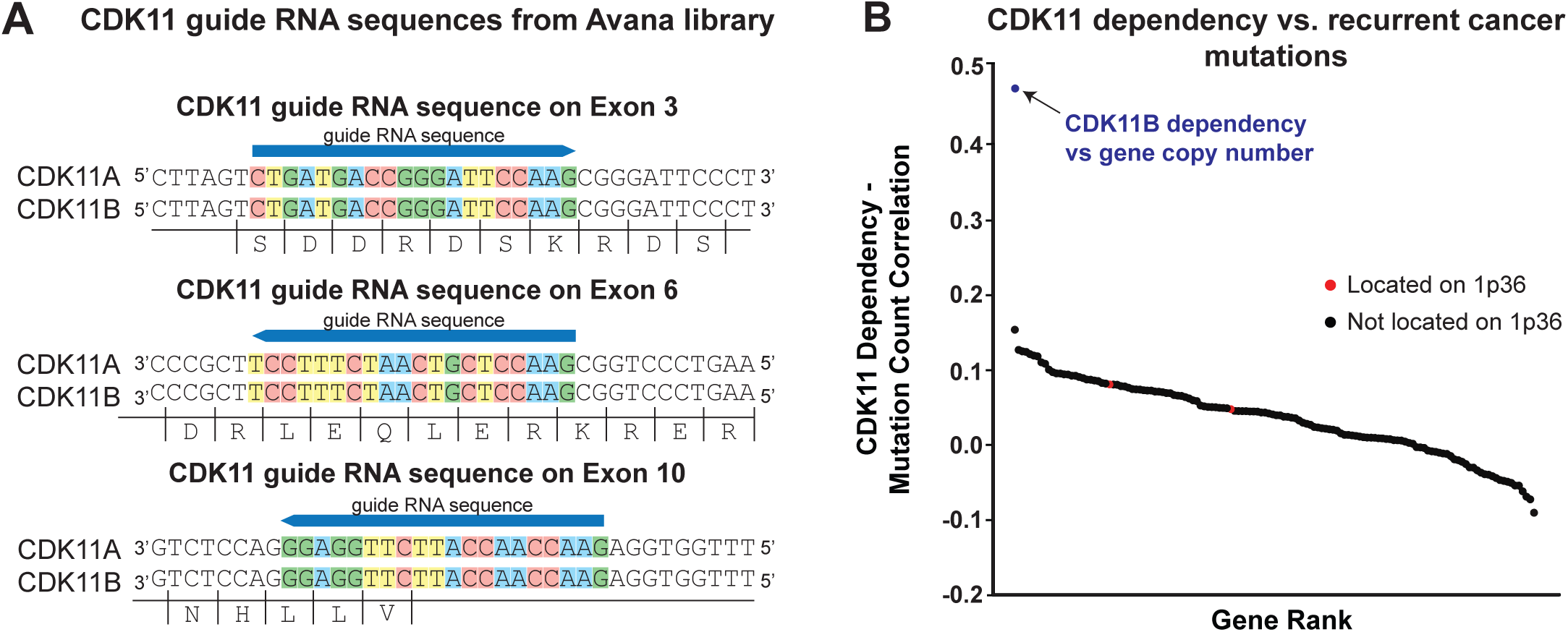
Analysis of CRISPR screening data targeting CDK11A/B. A) A schematic of guide RNA sequences from the Avana library that bind to both *CDK11A* and *CDK11B*. B) Waterfall plot displaying the correlation between recurrent cancer mutations and CDK11 dependency derived from CRISPR screening. Genes located on chromosome 1p36 are indicated in red. As a comparison, the correlation between CDK11 dependency and the copy number of *CDK11B* as calculated in Figure 4A is shown in blue. Complete results are presented in Table S9C.

**Supplemental Figure 7.**
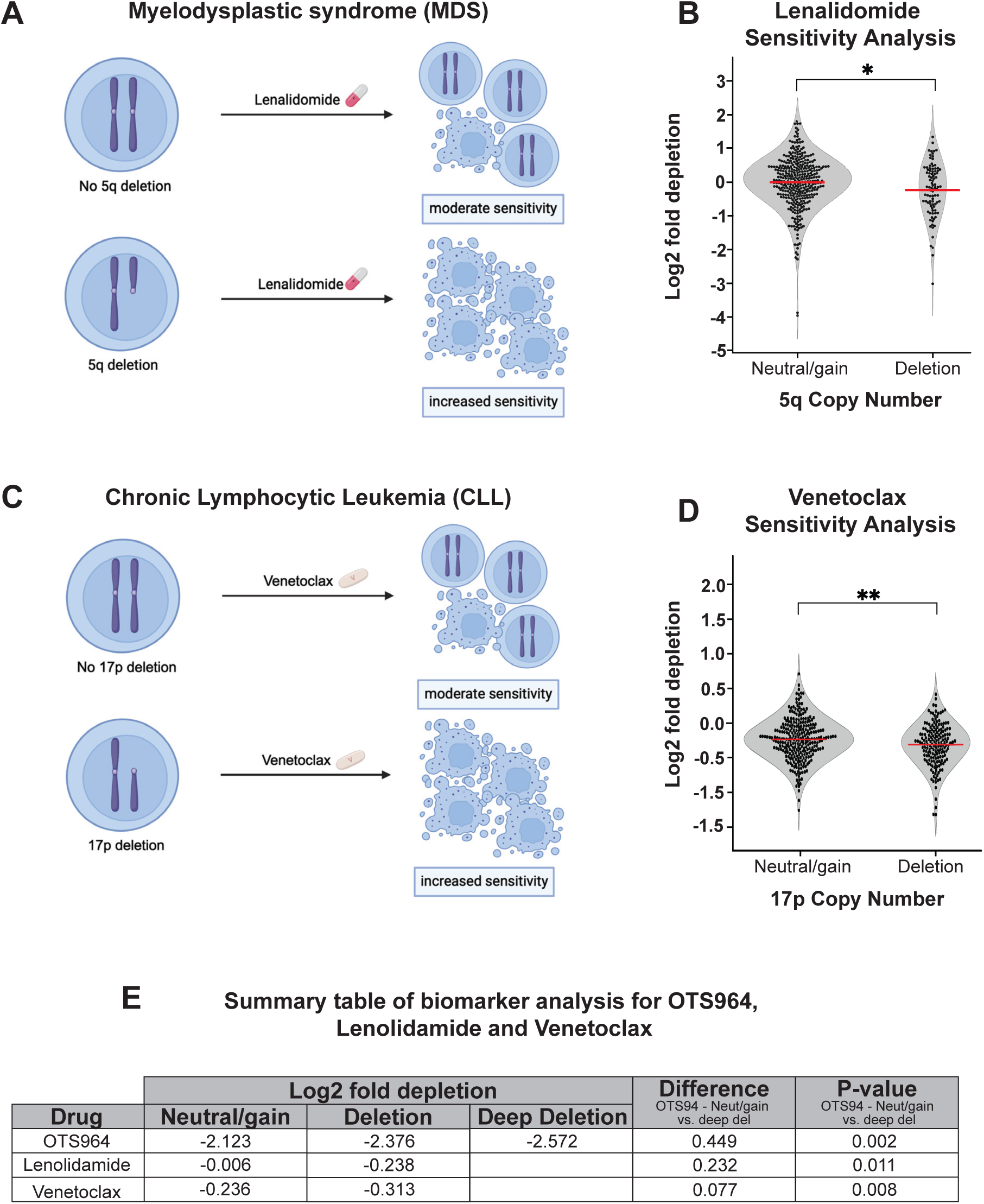
Assessing the association between chromosomal deletions and drug sensitivity for FDA-approved biomarkers. A) Schematic of 5q deletion as a biomarker for sensitivity to lenalidomide. B) Violin plots indicating the sensitivity of cancer cell lines to lenalidomide split based on the copy number of chromosome 5q. Every dot indicates a cancer cell line, and the red bars indicate the median of the data. The data is available via Corsello et al.^46^ C) Schematic of 17p deletion as a biomarker for sensitivity to venetoclax. D) Violin plots indicating the sensitivity of cancer cell lines to venetoclax split based on the copy number of chromosome 17p. Every dot indicates a cancer cell line, and the red bars indicate the median of the data. The data is available via Corsello et al.^46^

**Supplemental Figure 8.**
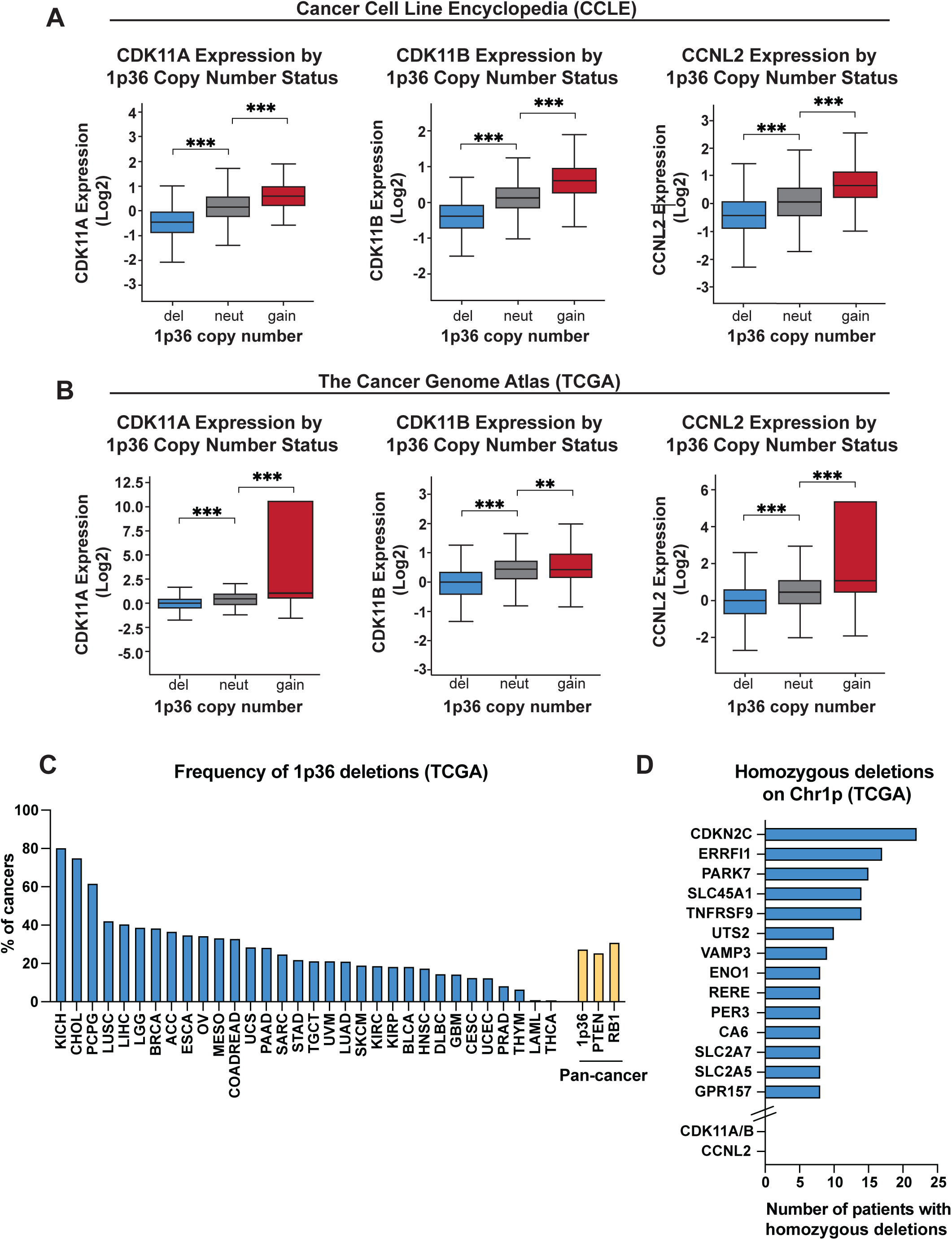
Heterozygous deletions of chromosome 1p36 are common across cancer types and are associated with decreased expression of CDK11 and CCNL2. A) Boxplots displaying the expression of *CDK11A* (left), *CDK11B* (middle), and *CCNL2* (right), split based on the copy number of chromosome 1p36 in cell lines from the Cancer Cell Line Encyclopedia. B) Boxplots displaying the expression of *CDK11A* (left), *CDK11B* (middle), and *CCNL2* (right), split based on the copy number of chromosome 1p36 in cancers from The Cancer Genome Atlas. C) Quantification of the frequency of chromosome 1p36 deletions in cancers from The Cancer Genome Atlas. As a comparison, we also analyzed the frequency of deletions in *PTEN* and *RB1*, shown in yellow. D) Quantification of the frequency of homozygous deletions in select genes encoded on chromosome 1p^90^. Complete results are presented in Table S11.

**Supplemental Figure 9.**
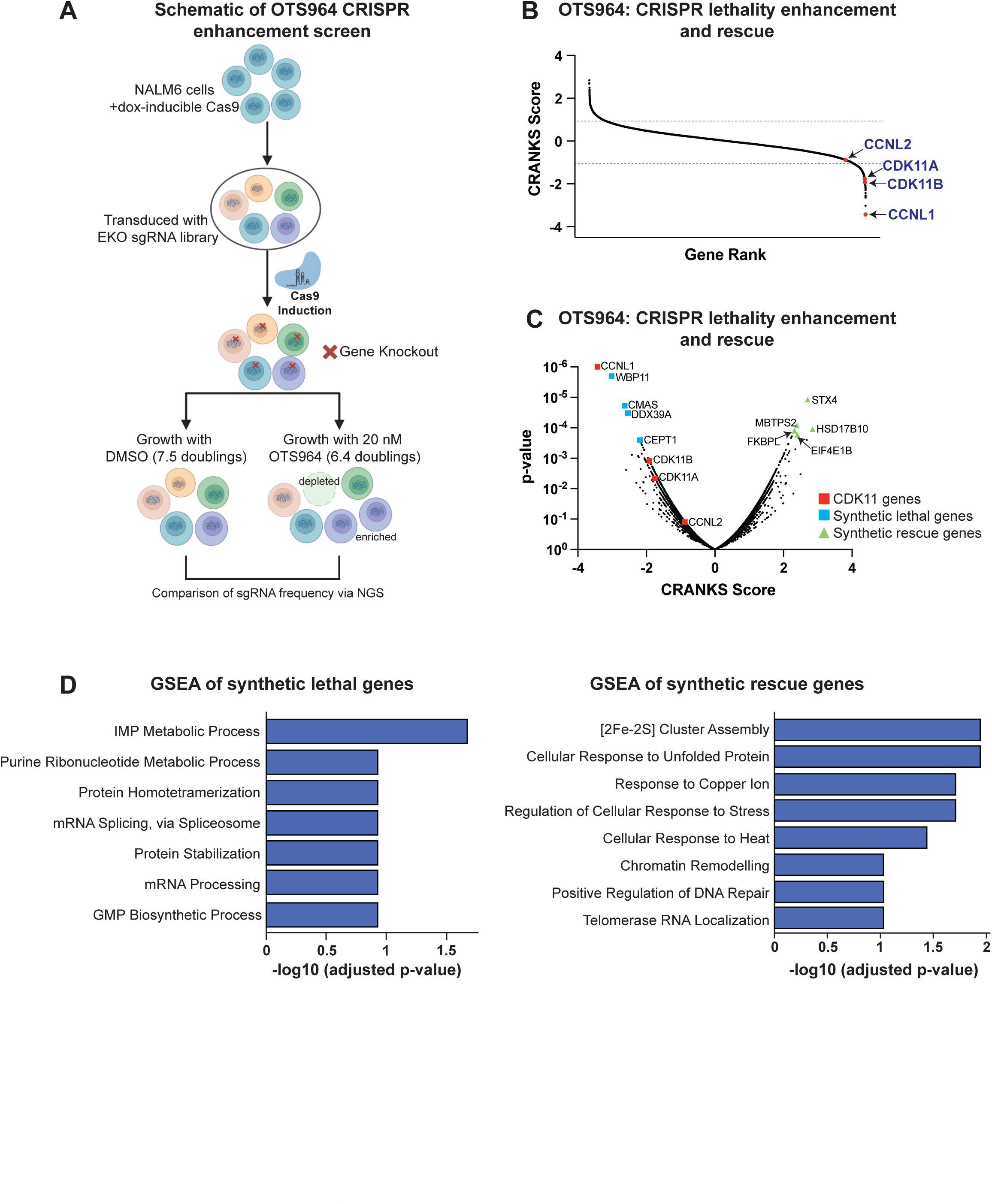
A genome-wide CRISPR screen to uncover modifiers of OTS964 sensitivity. A) Schematic of the CRISPR knockout screen performed in NALM6 cell lines treated with DMSO or 20 nM OTS964. B) Waterfall plot displaying the CRANKS Score obtained in CRISPR screen. CDK11 genes are highlighted in red. The dotted lines indicate synthetic lethality (-1) or synthetic rescue (1). Complete results are included in Table S13A. C) A volcano plot displaying CRANKS scores and p-values obtained from the CRISPR screen. CDK11 genes are highlighted in red. The top synthetic lethal genes are highlighted in blue and the top synthetic rescue genes are highlighted in green. D) A bar graph displaying GSEA results from the genes whose loss enhances sensitivity or resistance to OTS964. Complete results are included in Table S13B and S13C.

**Supplemental Figure 10.**
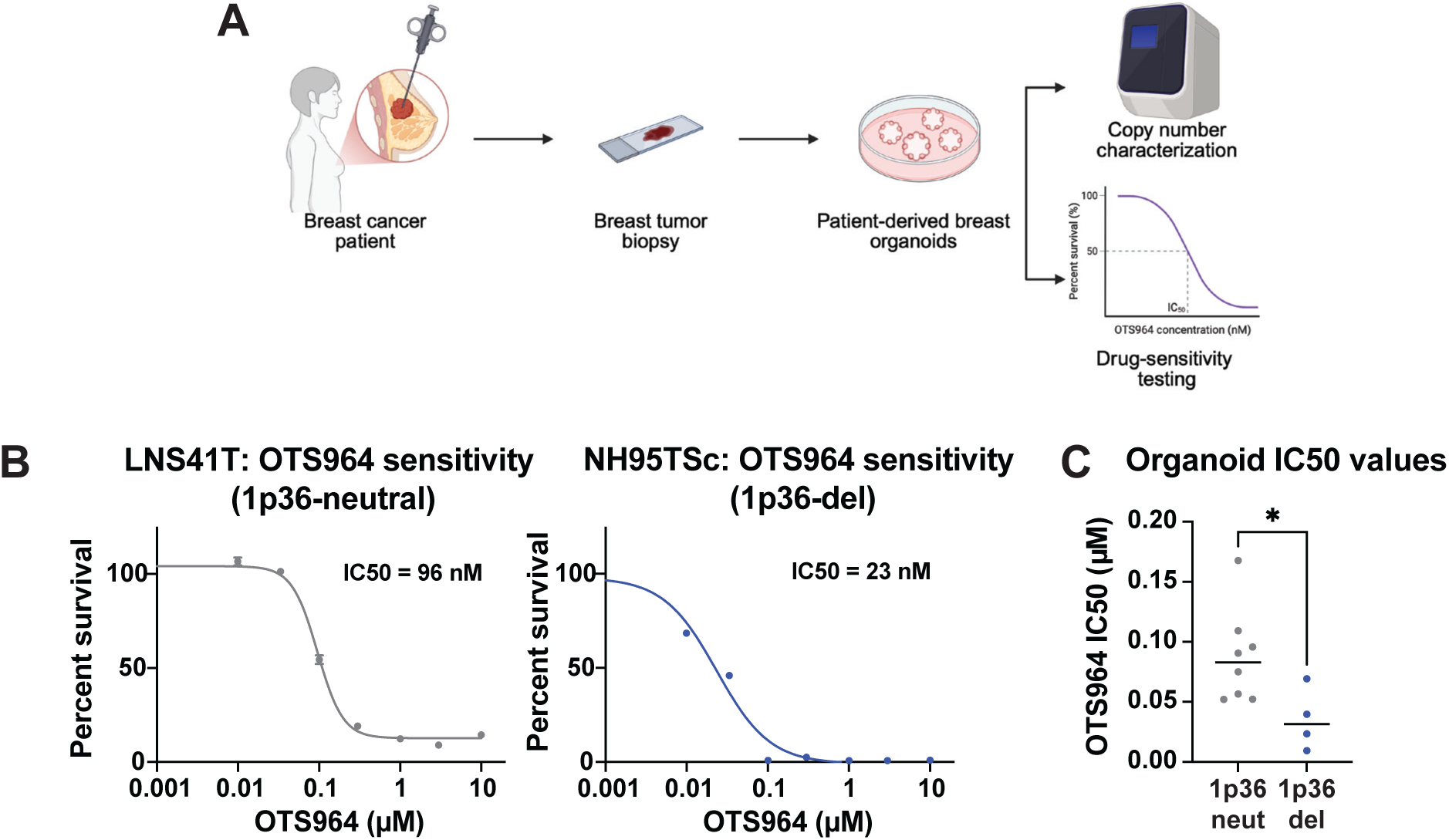
Chromosome 1p36 deletions are associated with increased sensitivity to OTS964 in patient-derived breast cancer organoids. A) Schematic of the collection and analysis of patient-derived organoids^50^. B) Dose-response curves showing survival of LNS4T1 (1p36-neutral) and NH95TSc (1p36-deletion) in response to varying concentrations of OTS964. The IC50 values for each cell line are annotated on the graph. C) Organoids treated with varying concentrations of OTS964. Graph displays the IC50 values comparing 1p36-neutral versus 1p36-deletion patient derived organoids. Statistical significance was determined by two-tailed t-tests (*, P < .05; **, P < .005; ***, P < .0005).

**Supplemental Figure 11.**
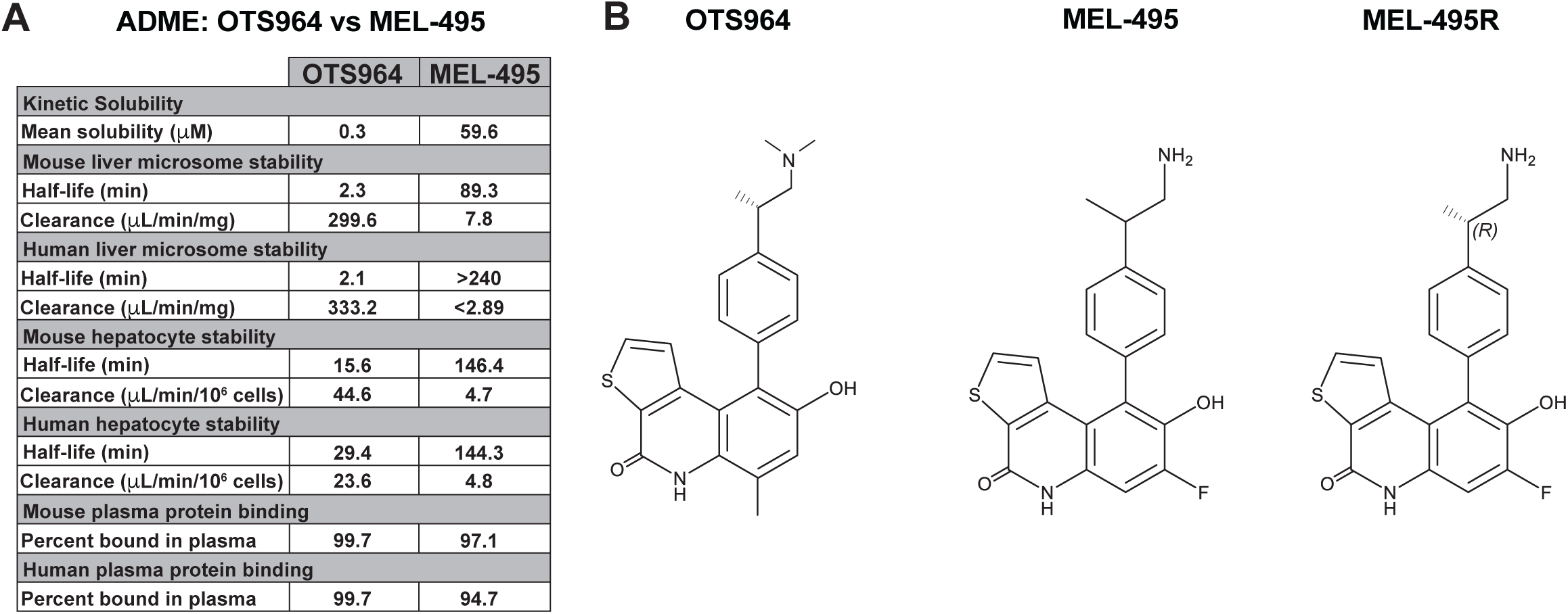
ADME analysis of OTS964, MEL-495, and MEL-495R. A) Table summarizing ADME profiles of OTS964 and MEL-495. B) Structures of the CDK11 inhibitors OTS964, MEL-495, MEL-495R.

**Supplemental Figure 12.**
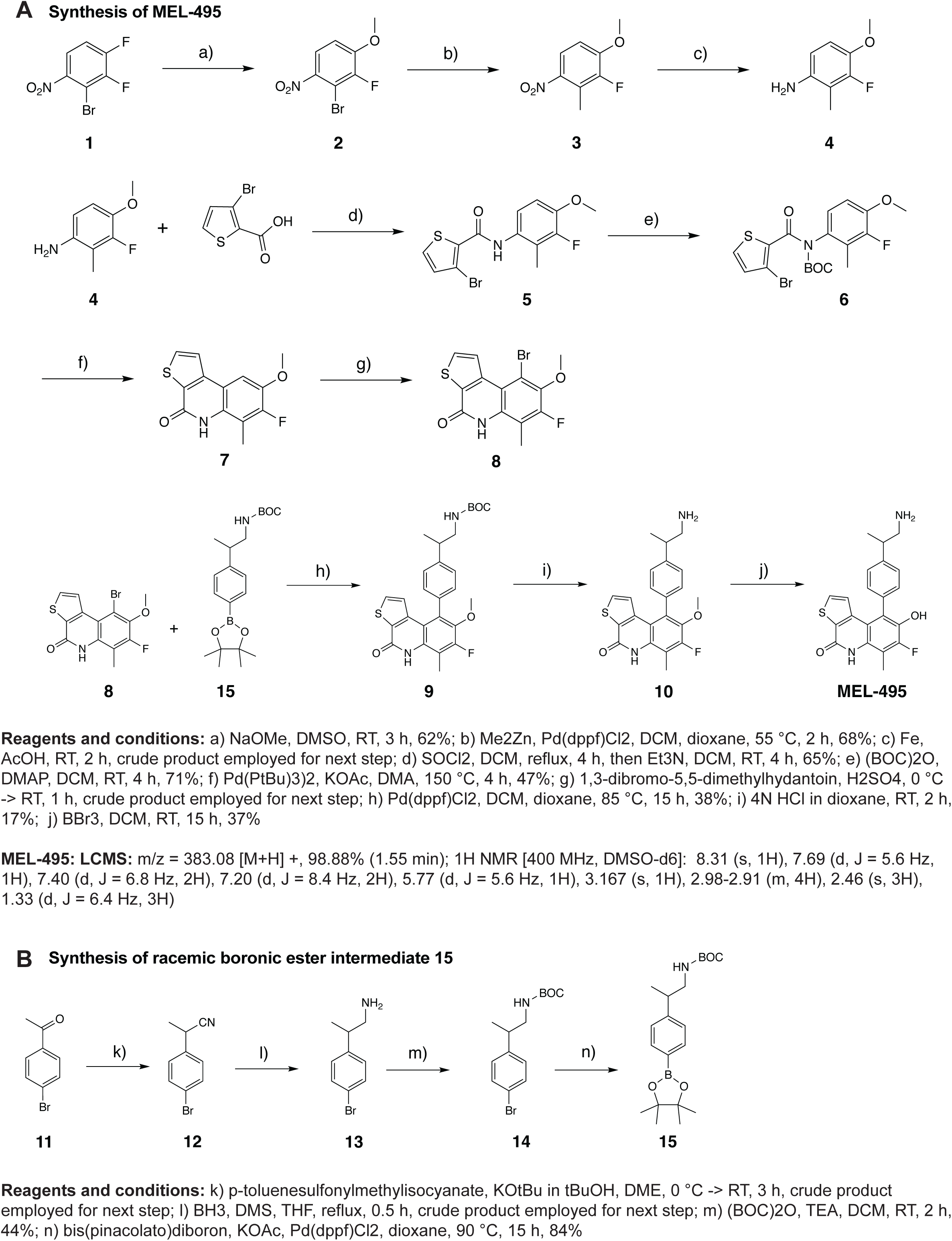
Synthesis of MEL-495. A schematic outlining the synthesis of MEL-495.

**Supplemental Figure 13.**
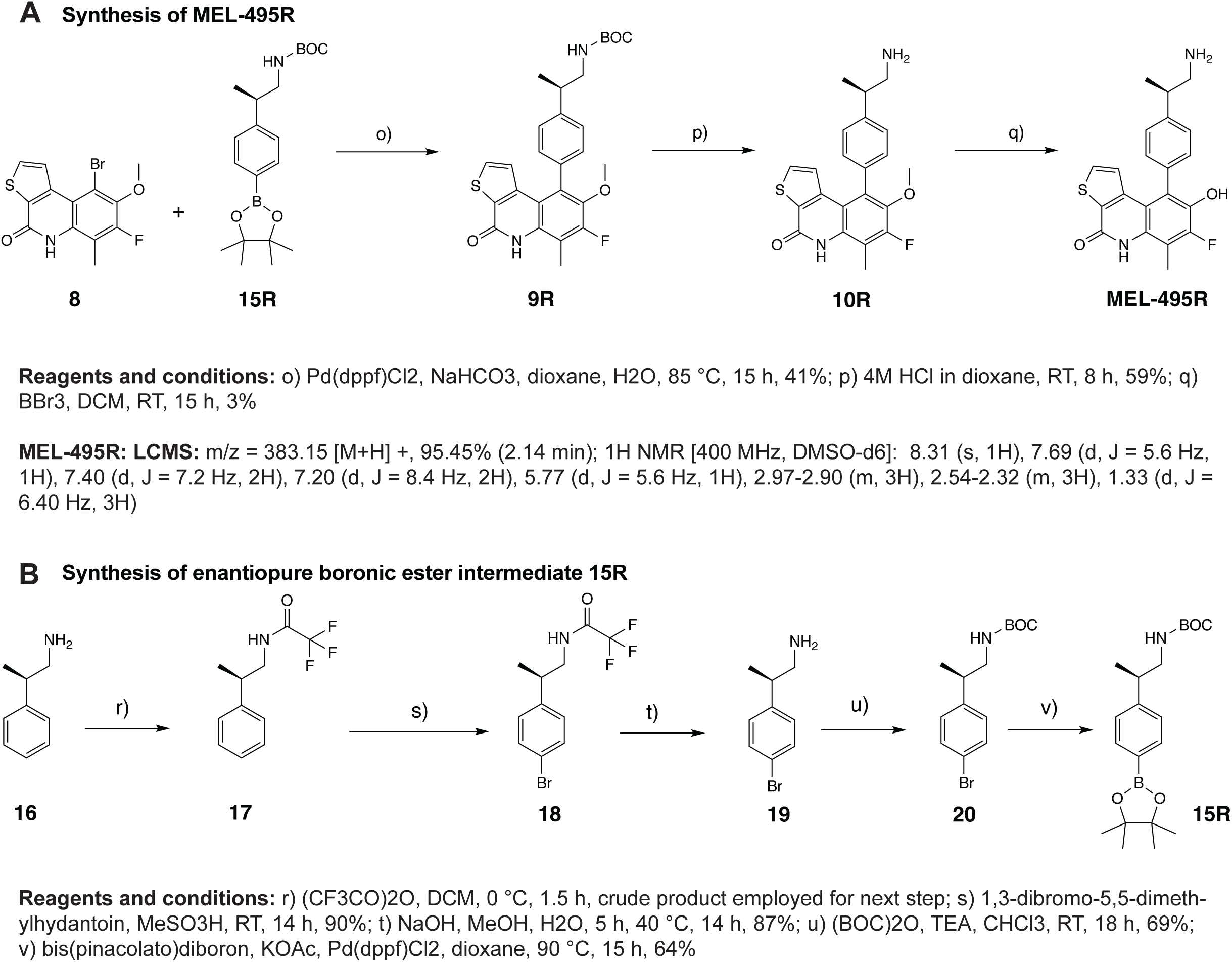
Synthesis of MEL-495R. A schematic outlining the synthesis of MEL-495R.

**Supplemental Figure 14.**
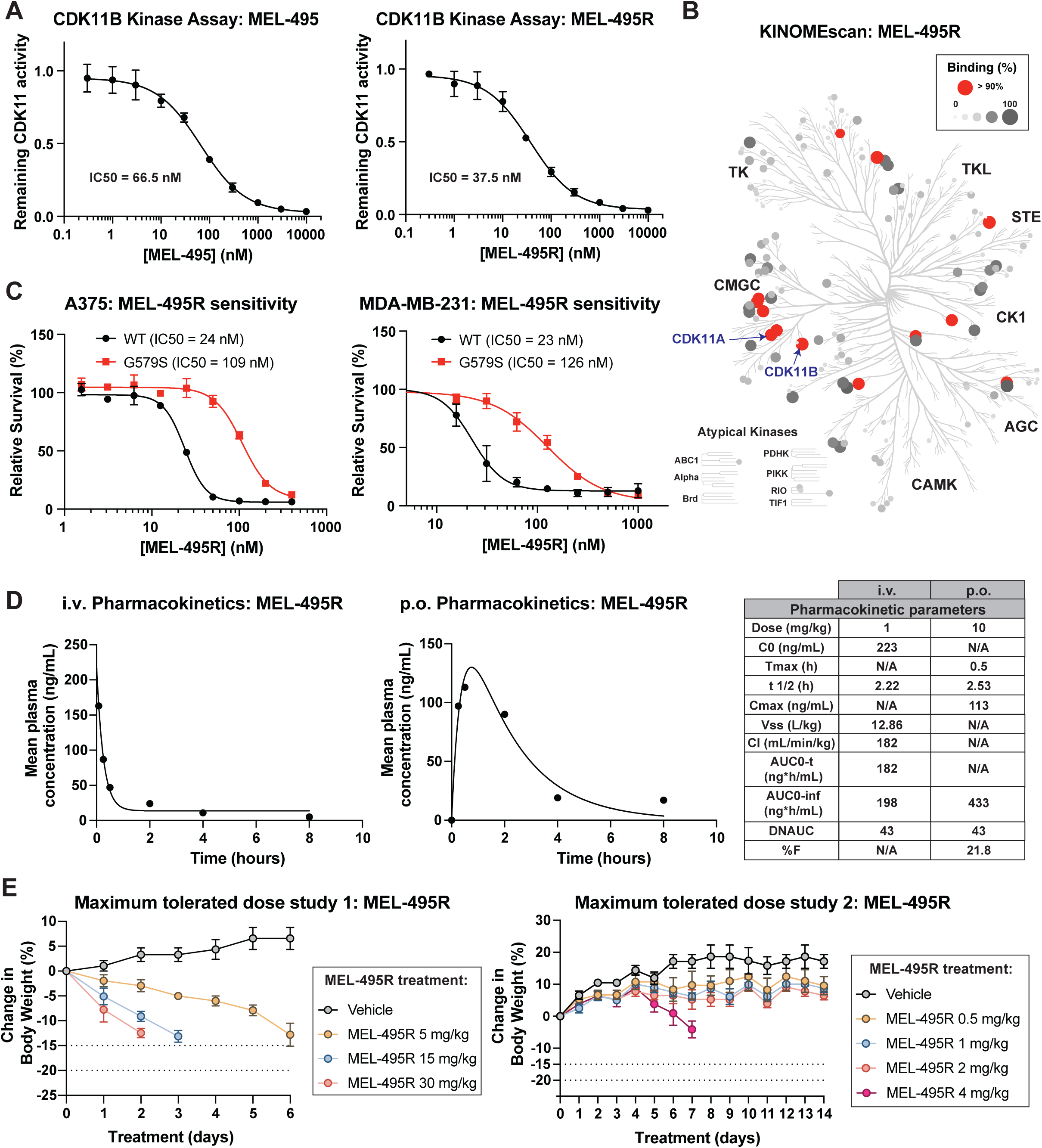
I*n vitro* and *in vivo* characterization of MEL-495R. A) CDK11B *in vitro* kinase activity assay. Dose response curves displaying CDK11B activity in the presence of either MEL-495 (left) or MEL-495R (right). B) KINOMEscan assay for 1 µM MEL-495R. The complete results are presented in Table S14. C) Dose-response curves showing the relative survival of A375 cells (left) and MDA-MB-231 cells (right) expressing wild-type CDK11B or CDK11B^G579S^ treated with varying concentrations of MEL-495R. D) Graphs showing the mean plasma concentration of MEL-495R after 1 mg/kg MEL-495R was delivered intravenously (left panel) or 10 mg/kg of MEL-495R was delivered orally (right panel). The table shows the pharmacokinetic parameters calculated for each route of administration. E) A maximum tolerated dose assay was performed for MEL-495R. The indicated dose of MEL-495R was delivered once daily via I.P. injection. Changes in body weight were monitored daily. Weight loss of 15% of the initial body weight of mouse was used as the endpoint of the study.

**Supplemental Figure 15.**
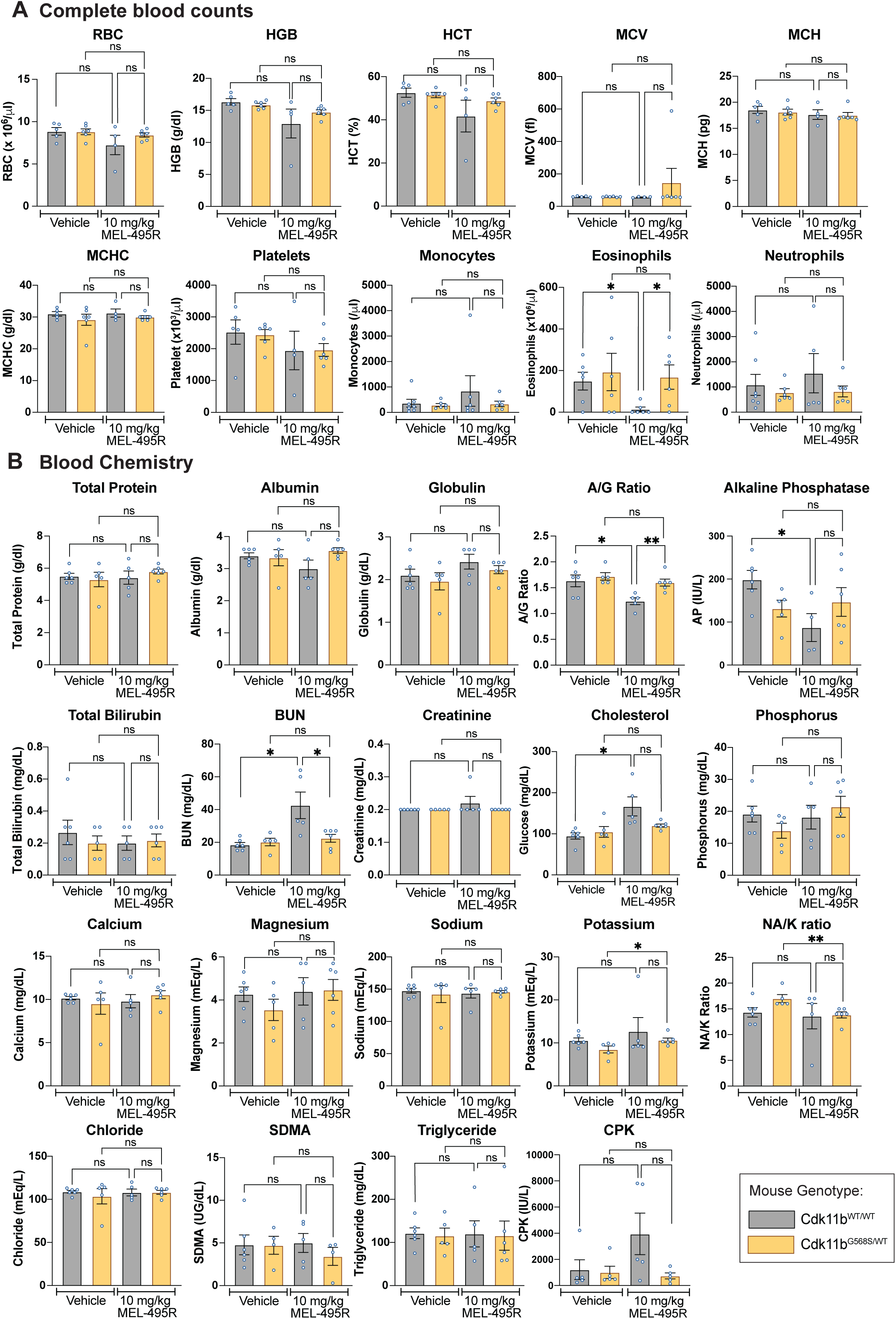
Additional blood cell counts and blood chemistry values in mice treated with MEL-495R. A) Full panel of complete blood counts as described in Figure 5G and 5I. Mean ± SEM of data from 4-5 mice per group. Complete data are presented in Table S15. Statistical significance was determined by unpaired t-tests (*, P < .05; **, P < .005; ***, P < .0005). B) Full panel of blood chemistry as described in Figure 5G and 5I. Complete data are presented in Table S15. Mean ± SEM of data from 4-5 mice per group. Statistical significance was determined by unpaired t-tests (*, P < .05; **, P < .005; ***, P < .0005).

**Supplemental Text 1: Confusion regarding the name CDK11.**

The protein now known as CDK11 has been referred to by multiple different names over time. CDK11 was first characterized as a 58 kilodalton protein with kinase activity, leading it to be initially called the p58 kinase^99^. Two facts about this kinase were quickly discovered: (1) it was homologous to the fission yeast CDC2 gene (which in humans is called CDK1) and (2) this protein was encoded by a pair of closely-related paralogs on human chromosome 1p36^57,100,101^. The kinase was said to be “CDC2-like”, leading these proteins to be named p58^CDC2L1^ and p58^CDC2L2^. Soon after this discovery, the kinase was renamed PITSLRE, based on a cyclin-binding amino acid motif that it encoded^57^. In the early 2000s, several groups independently uncovered the cyclin that binds to and activates this kinase^6,7,102^. As a result of this discovery, the p58/CDC2L1/PITSLRE kinase was renamed cyclin-dependent kinase 11, or CDK11.

However, at around the same time, a genome-wide survey of the human kinome was conducted^103^. This analysis identified a previously-unknown member of the CDK family encoded on chromosome 6q with strong homology to CDK8, which the authors also named CDK11. For several years after the publication of these competing reports, two entirely different kinases were both called CDK11.

Recognizing this confusion, Malumbres and colleagues proposed a uniform nomenclature for the human cyclin-dependent kinase family^104^. The original p58/CDC2L1/PITSLRE kinase was suggested to keep the name CDK11, while the CDK8-homolog discovered while characterizing the human kinome was renamed CDK19. Subsequently, the HUGO Gene Nomenclature Committee adopted these recommendations: CDK11A and CDK11B refer to the paralogous genes found on human chromosome 1p36 that were previously called p58, CDC2L1/2, and PITSLRE, while CDK19 encodes a homolog of CDK8 that is located on chromosome 6q^105–107^.

However, this complex history has led to lingering confusion and inaccuracies in the scientific literature and various genomic databases. For instance, in the Eurofins KINOMEscan assay, the protein now called CDK11B is called “CDC2L1” and the protein now called CDK19 is called “CDK19 (CDK11)”. Some recent papers also persist in using CDK11 to refer to CDK19. Finally, while CDK11 is conserved across unicellular and multicellular organisms, the names for its orthologs are not fully consistent: in *Drosophila melanogaster*, for example, the CDK11-like protein is still called PITSLRE^108^.

